# A red fluorescent genetically encoded biosensor for extracellular L-lactate

**DOI:** 10.1101/2022.08.30.505811

**Authors:** Yusuke Nasu, Yuki Kamijo, Rina Hashizume, Haruaki Sato, Yukiko Hori, Taisuke Tomita, Mikhail Drobizhev, Robert E. Campbell

## Abstract

L-Lactate, traditionally recognized as a waste product of metabolism, is now appreciated as a key intercellular energy currency in mammals. To enable investigations of intercellular shuttling of L-lactate, we have previously reported eLACCO1.1, a green fluorescent genetically encoded biosensor for extracellular L-lactate. eLACCO1.1 enables cellular resolution imaging of extracellular L-lactate in cultured mammalian cells and brain tissue. However, eLACCO1.1 spectrally overlaps with commonly used optical biosensors and actuators, limiting its application for multiplexed imaging or combined use with optogenetic actuators. Here, we report a red fluorescent extracellular L-lactate biosensor, designated R-eLACCO2. R-eLACCO2 is the end-product of extensive directed evolution and exhibits a large fluorescence response to L-lactate with high molecular specificity *in vitro*. We demonstrate that R-eLACCO2 with optimized leader and anchor sequences shows a large fluorescence change in response to extracellular L-lactate on the membrane of live mammalian cells. R-eLACCO2 should enable multicolor imaging of extracellular L-lactate in combination with other fluorescent probes and optogenetic actuators.

## Introduction

Living cells produce biologically-useful energy by catabolizing glucose to pyruvate which, in the presence of oxygen, is normally catabolized to carbon dioxide via the tricarboxylic acid (TCA) cycle or, in the absence of oxygen, is normally converted into L-lactate. One exception is aerobic glycolysis, or the so-called Warburg effect^1^, in which tumor cells produce L-lactate even under conditions where oxygen is available. Traditionally, L-lactate (the anionic form that predominates at pH > 3.9) and L-lactic acid (the protonated form that predominates at pH < 3.9) have been considered “waste” by-products of glucose metabolism^2^. In the early ninetieth century, Berzelius reported that the concentration of L-lactate was elevated in the muscles of hunted stags^3^. Much later, Fletcher and Hopkins showed that L-lactate accumulated in electrically-stimulated frog muscle under anaerobic condition^4^. In the 1970s, Fitts and Holloszy provided further support for the conclusions of these earlier studies by revealing the strong correlation between L-lactate concentration and muscle fatigue^5^. These landmark studies in the field of physiology led to the widely held view that L-lactate is the primary culprit determining muscle fatigue. The fact that a high level of L-lactate in serum is a clinical biomarker for acidosis, and associated with diseases such as sepsis and hypoxia, further reinforced the idea that L-lactate is a metabolic waste product^6^.

Recent studies have revealed that, contrary to its reputation as a waste product, L-lactate actually plays an important role as an intercellular and interorgan exchangeable fuel source in a range of tissues^7^. For example, circulating L-lactate rather than glucose can be a major source of fuel for the TCA cycle in all tissues of both fed and fasted mice^8^. Another example is the astrocyte-to-neuron lactate shuttle (ANLS) hypothesis which proposes that astrocytes metabolize glucose to produce L-lactate which is then released to the extracellular environment and taken up by neurons. In neurons, L-lactate is converted to pyruvate which is fed into the TCA cycle for production of adenosine triphosphate (ATP) to provide the energy necessary to sustain heightened neural activity^9,10^. In addition to the role of L-lactate as an energy fuel, there has been a growing appreciation of its role as a signaling molecule in a variety of tissues of mammals. Increased level of L-lactate released from cancer cells impairs cytokine production in tumor-infiltrating T cells and NK cells, thereby inhibiting tumor immunosurveillance and promoting tumor growth^11,12^. In neurons, astrocyte-derived L-lactate perturbs the intracellular redox state, hence stimulating the expression of genes related to neuronal plasticity, long-term memory and social behavior^9,10,13^. Disruption of L-lactate signaling between neurons and astrocytes is associated with diverse pathological conditions as Alzheimer’s disease, amyotrophic lateral sclerosis, depression, and schizophrenia^14,15^.

As extracellular L-lactate serves as both an energy fuel and a signaling molecule in numerous cellular processes, improved methods for monitoring of its concentration in live tissues are potentially of broad utility and high impact. To date, there have been several methods developed to monitor the concentration of extracellular L-lactate. One such method is the use of enzyme-based electrodes which use a probe inserted into the region of interest such as brain interstitium^16^. The concentration of extracellular L-lactate is determined *in situ* by enzymatic oxidation by immobilized lactate oxidase to produce hydrogen peroxide which is detected electrochemically. This method is applicable to *in vivo* L-lactate measurements, but the spatial resolution is limited and the insertion of the probe is invasive and causes local disruption of tissue. Several studies have demonstrated that pyruvate labeled with ^13^C isotope enables monitoring of L-lactate and other metabolites by using ^13^C-MRSI (magnetic resonance spectroscopic imaging)^17,18^. This method enables non-invasive monitoring of the level of extracellular L-lactate *in vivo*, but the spatiotemporal resolution is low and the sensitivity (detection limit) is not sufficient to detect sub-millimolar concentrations^19^. Commercially available kits can be utilized to estimate the concentration of L-lactate in cell media and biological fluids using the lactate oxidase catalyzed production of hydrogen peroxide which can be detected by colorimetric or fluorometric methods^17,20^. Such user-friendly methods are widely used, but cannot be practically adapted for continuous monitoring of the level of extracellular L-lactate *in situ* with high spatiotemporal resolution. Overall, existing methods are not sufficient to provide physiologically relevant information on the dynamics of extracellular L-lactate with cellular resolution.

A promising alternative to the methods described above would be a genetically encoded fluorescent biosensor based on the combination of a fluorescent protein and an L-lactate binding protein^21,22^. Introduction of the gene encoding such a fluorescent biosensor into a live organism could result in the expression of the biosensor protein in specific tissues of interest, allowing cell-based and *in vivo* fluorescence imaging of extracellular L-lactate with high spatiotemporal resolution. Such genetically encoded biosensors have revolutionized certain fields of biological science, most notably the field of neuroscience in which genetically encoded Ca^2+^ biosensors are widely used to monitor neural activity^23,24^.

Inspired by the tremendous impact that genetically encoded biosensors have had in various areas of biology and physiology, we and others have developed biosensors for the detection of intracellular^25−28^ and extracellular L-lactate^29^. For imaging of extracellular L-lactate, we recently reported a genetically encoded green fluorescent biosensor, designated eLACCO1.1, that was developed using an extensive directed evolution and an intensive effort to target it to the extracellular environment^29^. This first-in-class biosensor enables less-invasive imaging of extracellular L-lactate in cultured mammalian cells and brain tissue with cellular resolution, opening the door to a range of potential biological applications in which the dynamics of extracellular L-lactate are investigated in a cellular milieu.

Relative to green fluorescent biosensors such as eLACCO1.1, red fluorescent biosensors are associated with a number of inherent advantages, all other factors being the same. Relative to green light (∼500 nm), red light (∼600 nm) is inherently less phototoxic, is associated with reduced background autofluorescence, and exhibits greater tissue penetration, facilitating deeper and more sensitive imaging of tissues. In addition, red fluorescent biosensors enable multi-color imaging with blue and green biosensors and are compatible with the simultaneous use of blue light (∼450 nm) stimulated optogenetic actuators which can be used to control a range of cellular activities^30^. Since our lab reported the first genetically encoded red fluorescent Ca^2+^ biosensor R-GECO1 more than a decade ago, a variety of other red biosensors have been developed^22,31^. However, despite the remarkable progress in the development of such biosensors, relatively few genetically encoded red fluorescent biosensors for extracellular targets have yet been developed. One of the few examples is the red fluorescent R-iGluSnFR1 biosensor for neurotransmitter glutamate^32^. As additional examples, red fluorescent biosensors for extracellular neurotransmitter dopamine have been reported by two independent groups^33,34^. These examples demonstrate that red fluorescent biosensors can be targeted to, and functional in, the extracellular space. However, they tend to suffer from poor fluorescence response to target molecules compared to green fluorescent biosensors, hampering their wide applications in live tissues.

Here, we report the development of a genetically encoded red fluorescent biosensor for extracellular L-lactate. This biosensor, designated R-eLACCO2, is the end-product of extensive directed evolution and structure-based mutagenesis followed by optimization of biosensor expression and localization on the cell surface. We confirm that R-eLACCO2 has the highest fluorescence response of any genetically encoded red fluorescent biosensor in the extracellular environment, and enables imaging of extracellular L-lactate in cultured mammalian cells.

## Results

### Development of a genetically encoded red fluorescent L-lactate biosensor, R-eLACCO1.1

To construct an initial prototype red fluorescent L-lactate biosensor (**Fig. 1a**), we swapped the circularly permuted green fluorescent protein (cpGFP) of the green L-lactate biosensor eLACCO1 for the circularly permuted red fluorescent protein (cpmApple) (**Supplementary Fig. 1a**)^29^. The resulting variant, designated R-eLACCO0.1, showed a slight increase in fluorescence intensity (Δ*F*/*F* = (*F*_max_ - *F*_min_)/*F*_min_ = 0.2) upon treatment with L-lactate (**Supplementary Fig. 1b**). To develop variants of R-eLACCO0.1 with larger Δ*F*/*F*, we performed directed evolution with screening for L-lactate-dependent change in fluorescence intensity (**Fig. 1b**). Of a total of ten rounds of evolution, three rounds used a library in which both the N- and C-terminal linkers were randomized (**Supplementary Fig. 2**). The seven other rounds of evolution used libraries created by random mutagenesis of the entire biosensor gene. This effort ultimately produced the R-eLACCO1 variant with Δ*F*/*F* of 4.3 (**Fig. 1c** and **Supplementary Fig. 3a**).

**Fig. 1.**
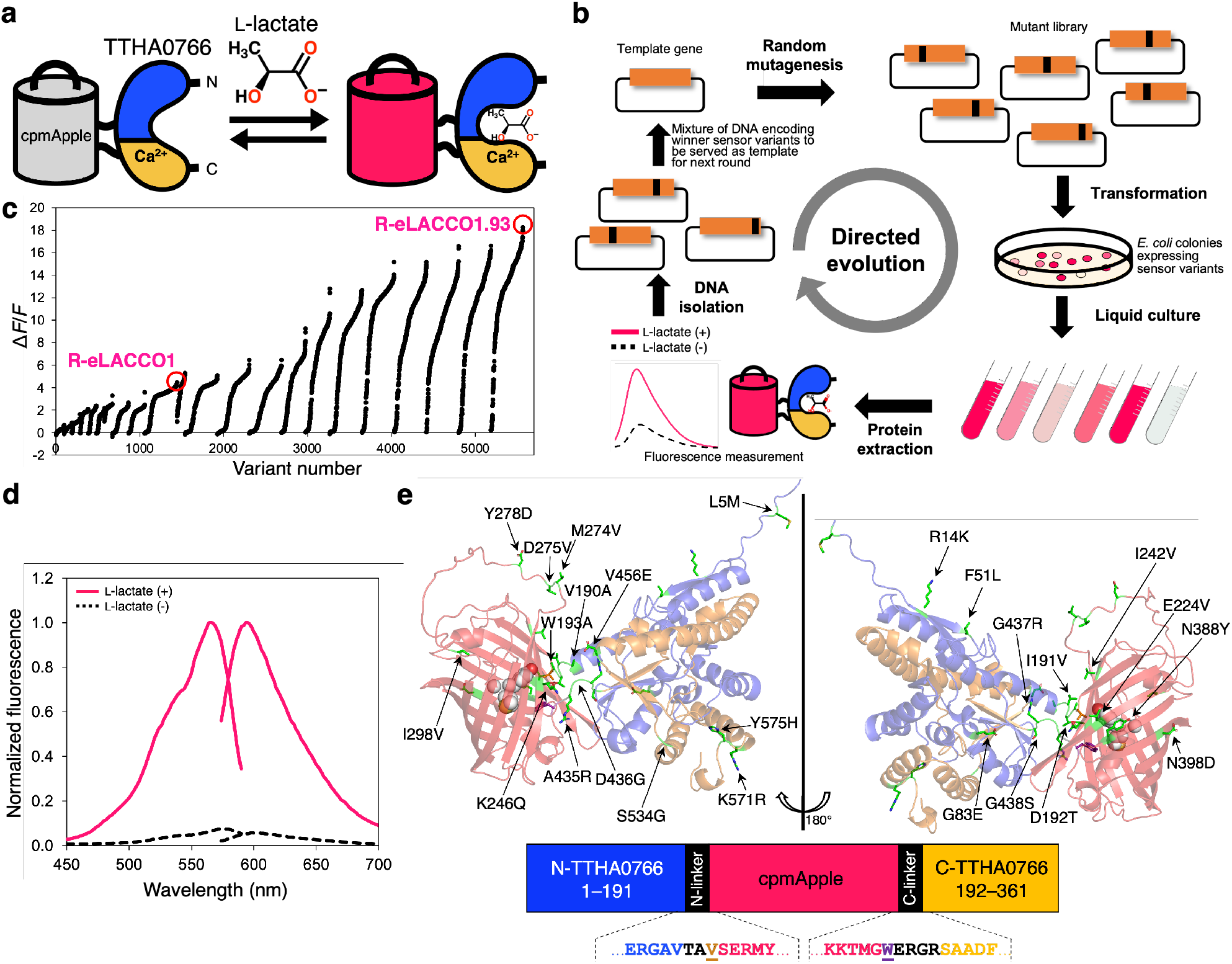
Development of R-eLACCO2. (**a**) Schematic representation of R-eLACCO and its mechanism of response to L-lactate. (**b**) Schematic of directed evolution workflow. Specific sites (i.e., the linkers) or the entire gene of template L-lactate biosensor were randomly mutated and the resulting mutant library was used to transform *E. coli*. Bright colonies were picked and cultured, and then proteins were extracted to examine Δ*F*/*F* upon addition of 10 mM L-lactate. Mixture of the variants with the highest Δ*F*/*F* were used as the template for the next round. (**c**) Δ*F*/*F* rank plot representing all proteins tested during the directed evolution. For each round, tested variants are ranked from lowest to highest Δ*F*/*F* value from left to right. (**d**) Excitation and emission spectra of R-eLACCO2 in the presence (10 mM) and absence of L-lactate. (**e**) Overall representation of the R-eLACCO2 model structure^47^ with the position of mutations indicated. The chromophore-forming tripeptide is shown in a sphere representation. In the primary structure of R-eLACCO2 (bottom), linker regions are shown in black and the two “gate post” residues^22^ in cpmApple are highlighted in dark orange (Val194) and purple (Trp433).

Green fluorescent extracellular L-lactate biosensor eLACCO1.1 has an apparent dissociation constant (*K*_d_) of 3.9 mM and shows a robust response to extracellular L-lactate when expressed on mammalian cells^29^. R-eLACCO1 has a 10× higher affinity (apparent *K*_d_ ∼ 310 μM) than eLACCO1.1 (**Supplementary Fig. 3b**). The high affinity of R-eLACCO1 inspired us to engineer a lower affinity variant of R-eLACCO1 for imaging of extracellular L-lactate. To tune the affinity of R-eLACCO1, we aimed to decrease its affinity for L-lactate using mutagenesis of residues that interact with L-lactate in the TTHA0766 lactate-binding domain. The crystal structure of eLACCO1 (PDB 7E9Y), which we assumed to be representative of TTHA0766 in R-eLACCO1, reveals that the phenol moiety of Tyr80 forms a hydrogen bond with bound L-lactate. To remove this hydrogen bond we introduced the Tyr80Phe mutation (**Supplementary Fig. 4a**), analogous to how we had previously converted the high affinity eLACCO1 to the low-affinity eLACCO1.1 (ref. 29). However, the resulting variant (R-eLACCO1 Tyr80Phe) exhibited a decreased fluorescence response to L-lactate, suggesting that Tyr80Phe mutation had substantially disrupted the L-lactate binding site (**Supplementary Fig. 4b**). Seeking an alternative approach to lower the affinity, we hypothesized that modification of a hydrophobic interaction could introduce a more modest change in the binding site and decrease the binding affinity. The crystal structure of eLACCO1 reveals that the hydrophobic side chain of Leu79 interacts with the methyl group of L-lactate (**Supplementary Fig. 4a**). Introduction of the conservative Leu79Ile mutation produced the low-affinity R-eLACCO1.1 variant with Δ*F*/*F* of 3.9 and apparent *K*_d_ = 1.4 mM (**Supplementary Fig. 5a,b**).

### *In vitro* characterization and cell-based imaging of R-eLACCO1.1

*In vitro* characterization revealed that R-eLACCO1.1 was a potentially useful biosensor for cell-based imaging of extracellular L-lactate (**Table 1** and **Supplementary Fig. 5a-g**). To explore the utility of R-eLACCO1.1, we expressed the protein targeted to the surface of mammalian cells and examined the fluorescence response to L-lactate (**Supplementary Fig. 5h**). Fluorescence imaging revealed that R-eLACCO1.1 is expressed on the cell surface, but the brightness is dim and the fluorescence response to L-lactate is limited (Δ*F*/*F* ∼ 1). The *in vitro* characterization had indicated that the molecular brightness of R-eLACCO1.1 is comparable to that of R-GECO1 cpmApple-based Ca^2+^ biosensor (**Table 1**), suggesting that the dim fluorescence in live mammalian cells is presumably due to the low expression level of R-eLACCO1.1. In addition, lactate-free R-eLACCO1.1 was found to harbor 75% of the neutral (dark) chromophore and 25% of the anionic (bright) chromophore, which changes to 55% neutral and 45% anionic for the L-lactate-bound state (**Table 1**). This chromophore equilibrium indicates that there could be substantial room to improve the Δ*F*/*F* by engineering decreased brightness of R-eLACCO1.1 in the lactate-free state and increased brightness of R-eLACCO1.1 in the lactate-bound state.

**Table 1.**
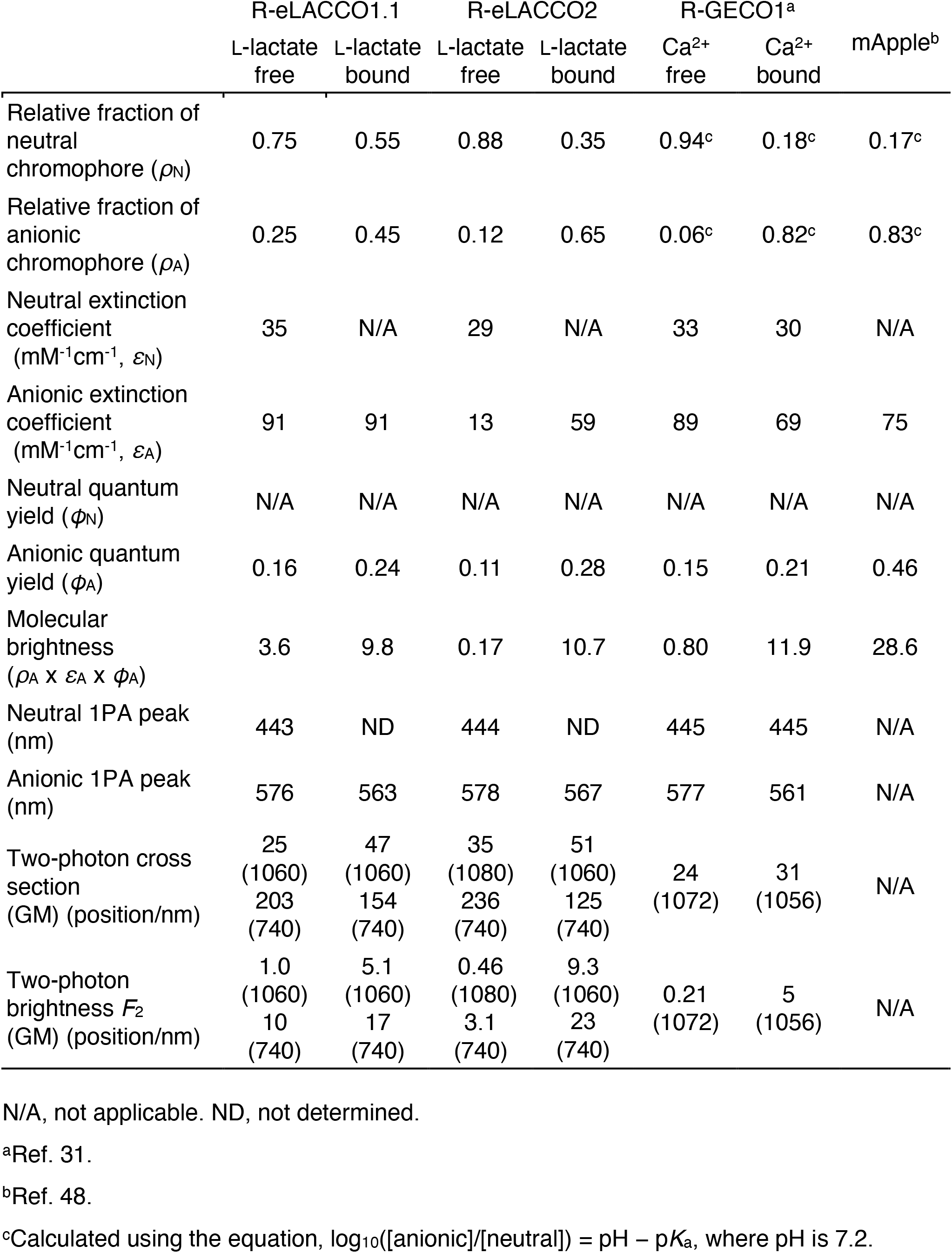
One- and two-photon photophysical parameters.

|  | R-eLACCO1.1 |  | R-eLACCO2 |  | R-GECO1<sup>a</sup> |  | mApple<sup>b</sup> |
| --- | --- | --- | --- | --- | --- | --- | --- |
|  | L-lactate<br>free | L-lactate<br>bound | L-lactate<br>free | L-lactate<br>bound | Ca<sup>2+</sup><br>free | Ca<sup>2+</sup><br>bound |  |
| Relative fraction of<br>neutral<br>chromophore ($\rho_N$) | 0.75 | 0.55 | 0.88 | 0.35 | 0.94<sup>c</sup> | 0.18<sup>c</sup> | 0.17<sup>c</sup> |
| Relative fraction of<br>anionic<br>chromophore ($\rho_A$) | 0.25 | 0.45 | 0.12 | 0.65 | 0.06<sup>c</sup> | 0.82<sup>c</sup> | 0.83<sup>c</sup> |
| Neutral extinction<br>coefficient<br>(mM<sup>-1</sup>cm<sup>-1</sup>, $\epsilon_N$) | 35 | N/A | 29 | N/A | 33 | 30 | N/A |
| Anionic extinction<br>coefficient<br>(mM<sup>-1</sup>cm<sup>-1</sup>, $\epsilon_A$) | 91 | 91 | 13 | 59 | 89 | 69 | 75 |
| Neutral quantum<br>yield ($\phi_N$) | N/A | N/A | N/A | N/A | N/A | N/A | N/A |
| Anionic quantum<br>yield ($\phi_A$) | 0.16 | 0.24 | 0.11 | 0.28 | 0.15 | 0.21 | 0.46 |
| Molecular<br>brightness<br>($\rho_A \times \epsilon_A \times \phi_A$) | 3.6 | 9.8 | 0.17 | 10.7 | 0.80 | 11.9 | 28.6 |
| Neutral 1PA peak<br>(nm) | 443 | ND | 444 | ND | 445 | 445 | N/A |
| Anionic 1PA peak<br>(nm) | 576 | 563 | 578 | 567 | 577 | 561 | N/A |
| Two-photon cross<br>section<br>(GM) (position/nm) | 25<br>(1060)<br>203<br>(740) | 47<br>(1060)<br>154<br>(740) | 35<br>(1080)<br>236<br>(740) | 51<br>(1060)<br>125<br>(740) | 24<br>(1072) | 31<br>(1056) | N/A |
| Two-photon<br>brightness $F_2$<br>(GM) (position/nm) | 1.0<br>(1060)<br>10<br>(740) | 5.1<br>(1060)<br>17<br>(740) | 0.46<br>(1080)<br>3.1<br>(740) | 9.3<br>(1060)<br>23<br>(740) | 0.21<br>(1072) | 5<br>(1056) | N/A |

N/A, not applicable. ND, not determined.
<sup>a</sup>Ref. 31.
<sup>b</sup>Ref. 48.
<sup>c</sup>Calculated using the equation, $\log_{10}([\text{anionic}]/[\text{neutral}]) = \text{pH} - \text{pK}_a$ , where pH is 7.2.

### Development of an improved biosensor, R-eLACCO2

To improve the expression level and Δ*F*/*F*, we further performed directed evolution using R-eLACCO1 as a starting template (**Fig. 1b,c** and **Supplementary Fig. 2**). One round of site-directed optimization, followed by eleven rounds of directed evolution by random mutagenesis of the whole gene, produced the high-performance variant R-eLACCO1.93 with Δ*F*/*F* of 18 (**Fig. 1c**). During this directed evolution process, we identified a beneficial mutation (Ile191Val) which is located next to N-terminal linker and is not included in R-eLACCO1.93. Introduction of this Ile191Val mutation to R-eLACCO1.93 ultimately led to a highly optimized variant, designated R-eLACCO2 (**Fig. 1d**). R-eLACCO2 contains a total of 25 mutations relative to R-eLACCO0.1 (**Fig. 1e** and **Supplementary Fig. 6**). Of the 25 mutations, eleven (L5M, R14K, F51L, G83E, V190A, I191V, G438S, V456E, S534G, K571R, and Y575H) are in TTHA0766, nine (E224V, I242V, K246Q, M274V, D275V, Y278D, I298V, N388Y, and N398D) are in cpmApple, and five (D192T, W193A, A435R, D436G, and G437R) are in the linkers. A non-responsive control biosensor, designated R-deLACCO1, was engineered by incorporating the Asp441Asn mutation into R-eLACCO2 to abolish L-lactate binding and subsequently performing six rounds of directed evolution to improve fluorescence brightness (**Supplementary Figs. 2** and **4c**).

### *In vitro* characterization of R-eLACCO2

With the highly-optimized R-eLACCO2 variant in hand, we undertook a detailed characterization of its biochemical and spectral properties. R-eLACCO2 displays an apparent *K*_d_ of 460 μM for L-lactate and Δ*F*/*F* of 20 upon treatment of 10 mM L-lactate (**Fig. 2a**). R-eLACCO2 has absorbance peaks at 444 nm and 578 nm, indicative of the neutral (protonated) and the anionic (deprotonated) chromophore, respectively, in the absence of L-lactate (**Fig. 2b**). Lactate binding results in the decrease in the neutral chromophore fraction and concomitant increase in the anionic chromophore fraction. Under one-photon excitation conditions, R-eLACCO2 in the absence of L-lactate displays an excitation peak at 574 nm and an emission peak at 598 nm. In the presence of L-lactate, these peak positions are slightly blue-shifted to 566 nm and 594 nm, respectively (**Fig. 1d**). The absorbance peak for the anionic form undergoes a similar blue-shift from 578 nm to 567 nm (**Fig. 2b**). The molecular brightness of R-eLACCO2 in the L-lactate bound state (excited at 563 nm which corresponds to the anionic form) is 90% of the R-GECO1 in Ca^2+^ bound state (**Table 1**)^31^. Bacterial colonies expressing R-eLACCO2 at 37 °C exhibited brighter fluorescence than R-eLACCO1 (**Fig. 2c**). Given the comparable L-lactate affinity and molecular brightness between R-eLACCO1 (*K*_d_ of 310 μM) and R-eLACCO2 (*K*_d_ of 460 μM), the increase in colony fluorescence suggests that R-eLACCO2 enhances the protein expression level at 37 °C relative to R-eLACCO1.

**Fig. 2.**
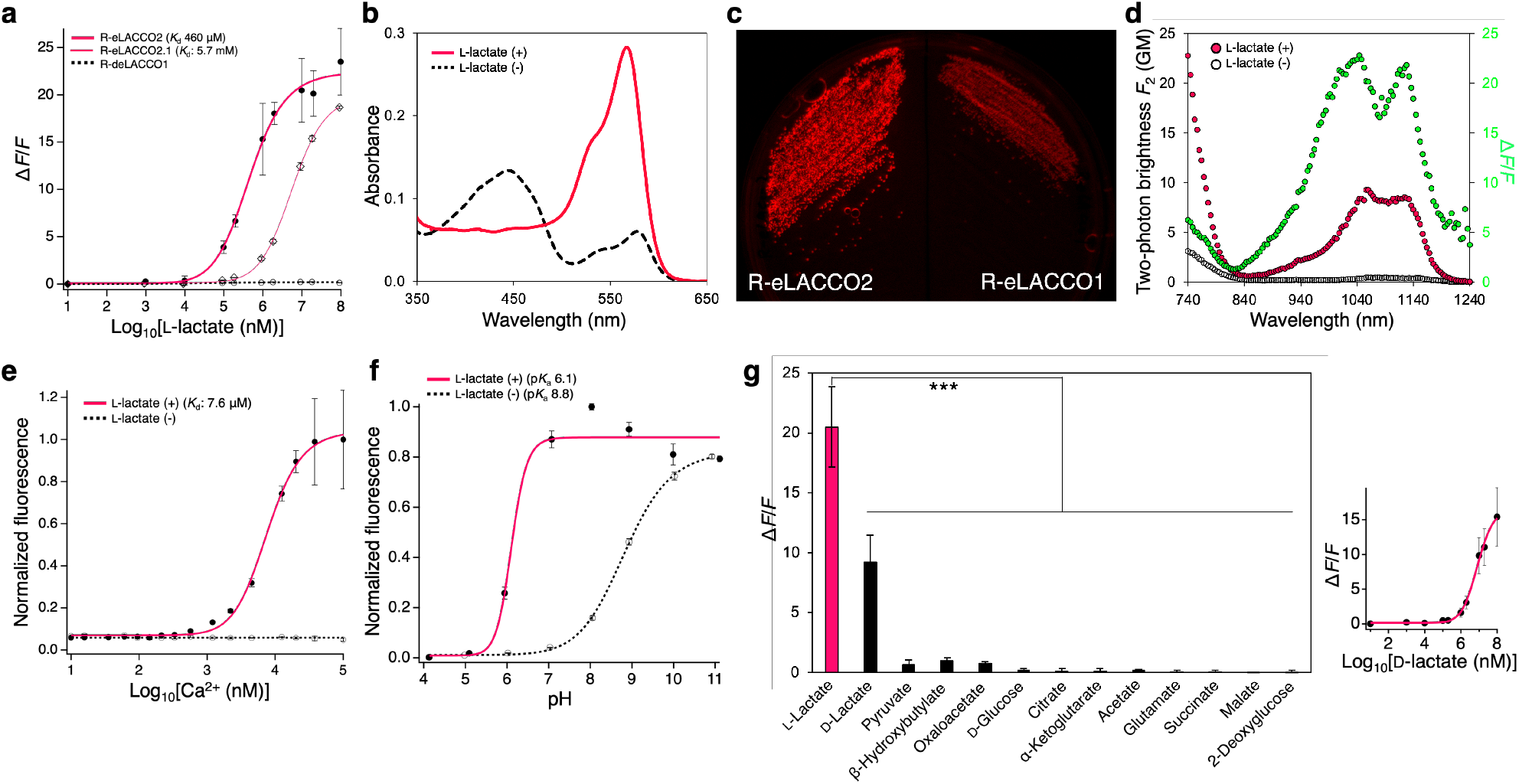
*In vitro* characterization of R-eLACCO2. (**a**) Dose-response curves of R-eLACCO2, R-eLACCO2.1, and R-deLACCO1 for L-lactate. *n* = 3 experimental triplicates (mean ± s.d.). (**b**) Absorbance spectra of R-eLACCO2 in the presence (50 mM) and absence of L-lactate. (**c**) Fluorescence image of a streak plate of *E. coli* expressing R-eLACCO1 or R-eLACCO2. (**d**) Two-photon excitation spectra of R-eLACCO2 in the presence (10 mM) and absence of L-lactate. Δ*F*/*F* is represented in the green plots. GM, Goeppert-Mayer units. (**e**) Dose-response curve of R-eLACCO2 as a function of Ca^2+^ in the presence (100 mM) and absence of L-lactate. *n* = 3 experimental triplicates (mean ± s.d.). (**f**) pH titration curves of R-eLACCO2 in the presence (100 mM) and absence of L-lactate. *n* = 3 experimental triplicates (mean ± s.d.). (**g**) Specificity of R-eLACCO2. Concentration of each metabolite is 10 mM. Right graph represents a dose-response curve of R-eLACCO2 for D-lactate. *n* = 3 experimental triplicates (mean ± s.d.). Statistical analysis was performed using one-way analysis of variance (ANOVA) with the Dunnett’s post hoc tests. \*\*\**p* < 0.000001.

The two-photon absorption spectra of R-eLACCO2 show a double-humped shape in both the presence and absence of L-lactate, similar to those of R-GECO1 (ref. 35). R-eLACCO2 in the absence of L-lactate displays peaks at 1076 nm and 1148 nm, while it exhibits peaks at 1060 nm and 1124 nm in the presence of L-lactate (**Fig. 2d**). This result indicates that R-eLACCO2 blue shifts upon L-lactate binding under two-photon excitation, consistent with one-photon excitation. The intrinsic two-photon brightness is calculated as *F*_2_ = σ_2,A_ × φ_A_ × ρ_A_ + σ_2,N_ × φ_N_ × ρ_N_, where indices A and N correspond to the anionic and neutral forms, σ_2_ is the two-photon absorption cross section, φis the fluorescence quantum yield, and ρ is the relative fraction of neutral or anionic form. The brightness of L-lactate-bound R-eLACCO2 excited at 1060 nm, *F*_2_ = 9.3 GM, is larger than that of Ca^2+^-bound R-GECO1 (*F*_2_ = 5 GM) (**Table 1**)^35^. The L-lactate induced two-photon excited fluorescence change (11*F*_2_/*F*_2_ = 20 at 1060 nm) is comparable to the one-photon excited fluorescence change (11*F*_1_/*F*_1_ = 20 at 593 nm) (**Figs. 1d** and **2d**).

The crystal structure of eLACCO1 had revealed a critical Ca^2+^ in the L-lactate binding pocket^29^. To investigate the effect of Ca^2+^ on R-eLACCO2 functionality, we measured the fluorescence intensity in a range of Ca^2+^ concentration. These experiments revealed that Ca^2+^ is indeed essential for the L-lactate-dependent response of R-eLACCO2, and the biosensor only functions as an L-lactate biosensor at concentrations of Ca^2+^ greater than ∼38 μM (**Fig. 2e**). R-eLACCO2 showed a pH dependence with p*K*_a_ values of 6.1 and 8.8 in the presence and absence of L-lactate, respectively, similar to that of R-GECO1 (**Fig. 2f** and **Table S1**)^31^. R-deLACCO1 showed no response to L-lactate (**Fig. 2a**), and pH dependence that was similar to the lactate-free state of R-eLACCO2 (**Supplementary Fig. 7c**).

Investigation of the molecular specificity revealed that R-eLACCO2 is highly specific for L-lactate over a wide array of metabolites (**Fig. 2g**). R-eLACCO2 responds to D-lactate with an apparent *K*_d_ of 8.4 mM, a concentration that is far greater than the ∼11–70 nM concentration in serum^36^. Overall, these results indicate that R-eLACCO2 responds only to changes in L-lactate concentration and pH under physiological conditions.

### Development of R-eLACCO2.1 and targeting to the extracellular environment

Introduction of the Leu79Ile mutation to R-eLACCO2 produced a low-affinity R-eLACCO2.1 variant with apparent *K*_d_ of 5.7 mM (**Fig. 2a** and **Supplementary Fig. 7**). Our previous work on eLACCO1.1 had demonstrated that the combination of N-terminal leader sequence and C-terminal anchor domain plays an essential role in the performance of biosensor for extracellular target^29^. To identify the best leader-anchor combination, we first examined the efficiency of cell surface targeting of R-eLACCO2.1 variants with human CD59-derived leader sequence and a range of anchor domains (**Fig. 3a,b**). Colocalization analyses of these R-eLACCO2.1 variants and a green fluorescent cell surface marker revealed that all protein-based anchors, including a widely-used platelet-derived growth factor receptor (PDGFR), resulted in only intracellular expression with localization reminiscent of the nuclear membrane or Golgi apparatus. In contrast, all lipid (glycosylphosphatidylinositol, GPI)-based anchors resulted in efficient targeting to cell surface. Three (CD59, COBRA, and GFRA1) of the four lipid-based anchors investigated resulted in a high proportion of biosensor localized on the cell surface.

**Fig. 3.**
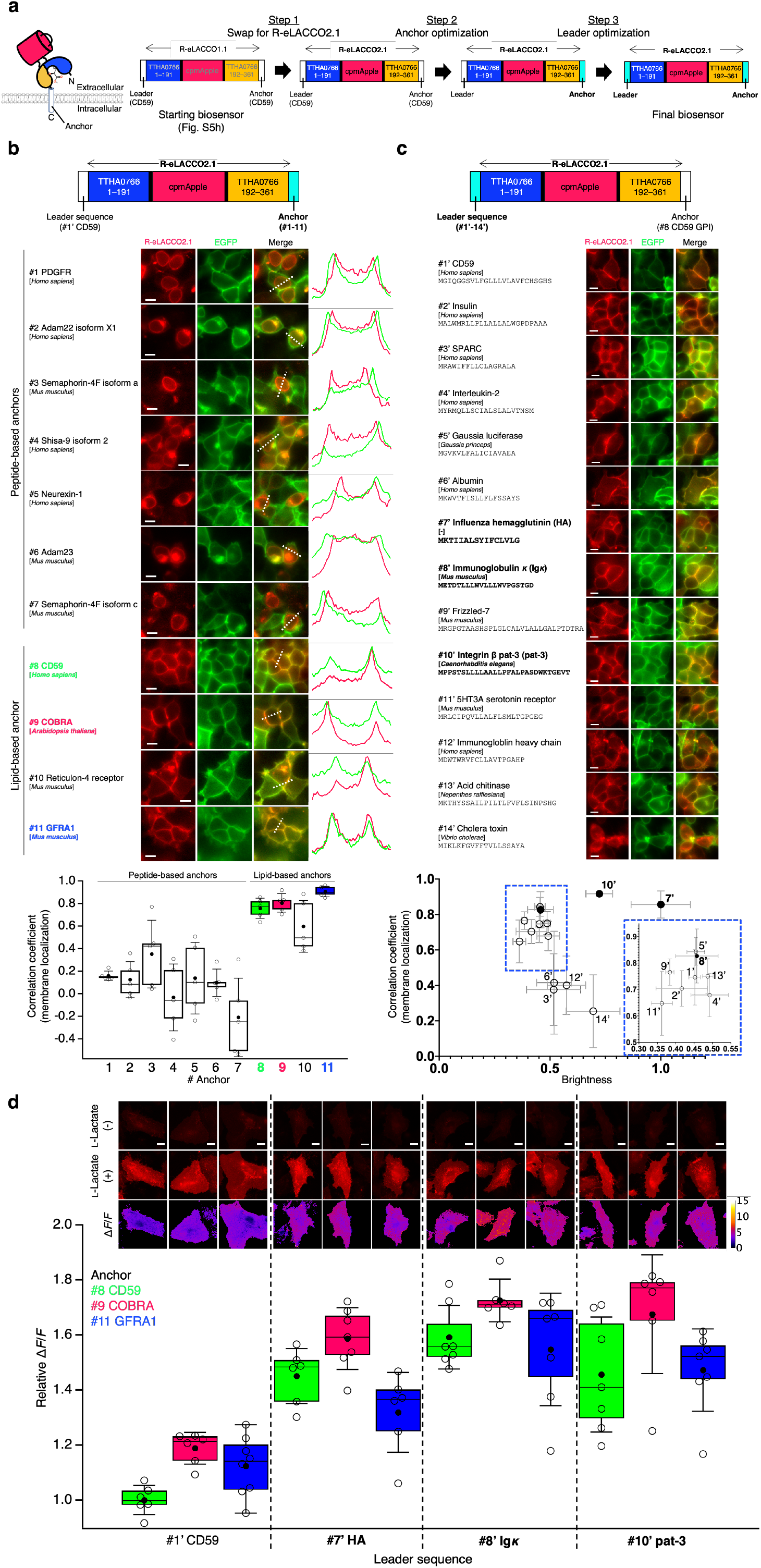
Optimization of targeting of R-eLACCO2.1 to the extracellular membrane surface. (**a**) Overview of the biosensor optimization. (**b**) Localization of R-eLACCO2.1 with CD59 leader and a range of anchors in HEK293T cells. Line-scans (right) correspond to dashed white lines on the merged images. EGFP with CD59 leader and CD59 GPI anchor was used as cell surface marker. *n* = 5 cells per construct for correlation coefficient. Scale bars, 10 μm. (**c**) Localization of R-eLACCO2.1 with various leader sequences and CD59 GPI anchor in HEK293T cells. EGFP with CD59 leader and CD59 GPI anchor was used as cell surface marker. Mean ± s.e.m., *n* = 4 cells per construct for correlation coefficient, *n* = 4 field of views (FOVs) per construct for brightness. Scale bars, 10 μm. (**d**) Relative Δ*F*/*F* of R-eLACCO2.1 with a range of combinations of leader and anchor in HeLa cells. Scale bars, 20 μm. In the box plots of (b) and (d), the horizontal line is the median; the top and bottom horizontal lines are the 25^th^ and 75^th^ percentiles for the data; and the whiskers extend one standard deviation range from the mean represented as black filled circle.

We next screened various N-terminal leader sequences in combination with the CD59 GPI anchor (**Fig. 3c**). Of 14 leader sequences examined, two (SPARC and cholera toxin) resulted in only intracellular localization, while the other twelve gave excellent membrane localization of R-eLACCO2.1. This result suggests that both the C-terminal anchor and the N-terminal leader sequence can influence targeting to the cell surface. Three leader sequences (influenza hemagglutinin (HA), immunoglobulin *κ* (Ig*κ*), and integrin β pat-3 (pat-3)) provided the brightest fluorescent signals and were therefore chosen as the most promising leader sequences. To determine the best combination of the leader sequences and anchor domains, we combined the most promising leader sequences (HA, Ig*κ*, and pat-3) and anchor domains (CD59, COBRA, and GFRA1) and measured Δ*F*/*F* of each combination in HeLa cells upon treatment of 10 mM L-lactate (**Fig. 3d**). This measurement revealed that the best combination for cell surface targeting of R-eLACCO2.1 is the Ig*κ*-derived N-terminal leader sequence and the COBRA-based C-terminal anchor. Highly optimized R-eLACCO2.1 with the best leader-anchor combination shows Δ*F*/*F* of ∼5 which is approximately 5× higher than R-eLACCO1.1 with the CD59-derived leader and anchor combination (**Supplementary Fig. 5h**).

### Characterization of R-eLACCO2 variants in live mammalian cells

We characterized R-eLACCO2 (high affinity variant) and R-eLACCO2.1 (low affinity variant) variants with the optimized leader (Ig*κ*) and anchor (COBRA) expressed in mammalian cells (**Fig. 4a**). The application of 10 mM L-lactate robustly increased fluorescence intensity of R-eLACCO2 (Δ*F*/*F* of 4.5 ± 0.1, mean ± s.e.m.) and R-eLACCO2.1 (Δ*F*/*F* of 6.2 ± 0.2, mean ± s.e.m.) expressed on HeLa cells (**Fig. 4b,c**). The control biosensor R-deLACCO1 had similar membrane localization and, as expected, did not respond to L-lactate even at the highest concentration tested (**Fig. 4a-d**). R-eLACCO2 and R-eLACCO2.1 have *in situ* apparent *K*_d_s of 560 μM and 5.1 mM for L-lactate, respectively (**Fig. 4d**). Similar to TTHA0766-based eLACCO1.1 (ref. 29), R-eLACCO2 and R-eLACCO2.1 display Ca^2+^ dependent fluorescence with apparent *K*_d_s of 310 μM and 610 μM, respectively (**Fig. 4e**). These *in situ K*_d_s, which are higher than those measured with purified proteins (**Fig. 2e** and **Supplementary Fig. 7b**), are much lower than the extracellular Ca^2+^ concentration (1.5−1.7 mM) in brain^37^. To probe the response kinetics of R-eLACCO2 variants, we bathed R-eLACCO-expressing HeLa cells in a solution containing 0 mM and 10 mM L-lactate. The bath application imaging revealed that R-eLACCO variants had a slightly faster on rate (τ_on_ of 61 ± 5 s for R-eLACCO2 and 61 ± 3 s for R-eLACCO2.1, mean ± s.d.) than the green fluorescent eLACCO1.1 (τ_on_ of 68 ± 8 s, mean ± s.d.) (**Fig. 4f**). The off rates of R-eLACCO2 (τ_off_ of 72 ± 4 s, mean ± s.d.) and R-eLACCO2.1 (τ_off_ of 35 ± 3 s, mean ± s.d.) are slower than that of eLACCO1.1 (τ_off_ of 22 ± 5 s, mean ± s.d.) (**Fig. 4f**).

**Fig. 4.**
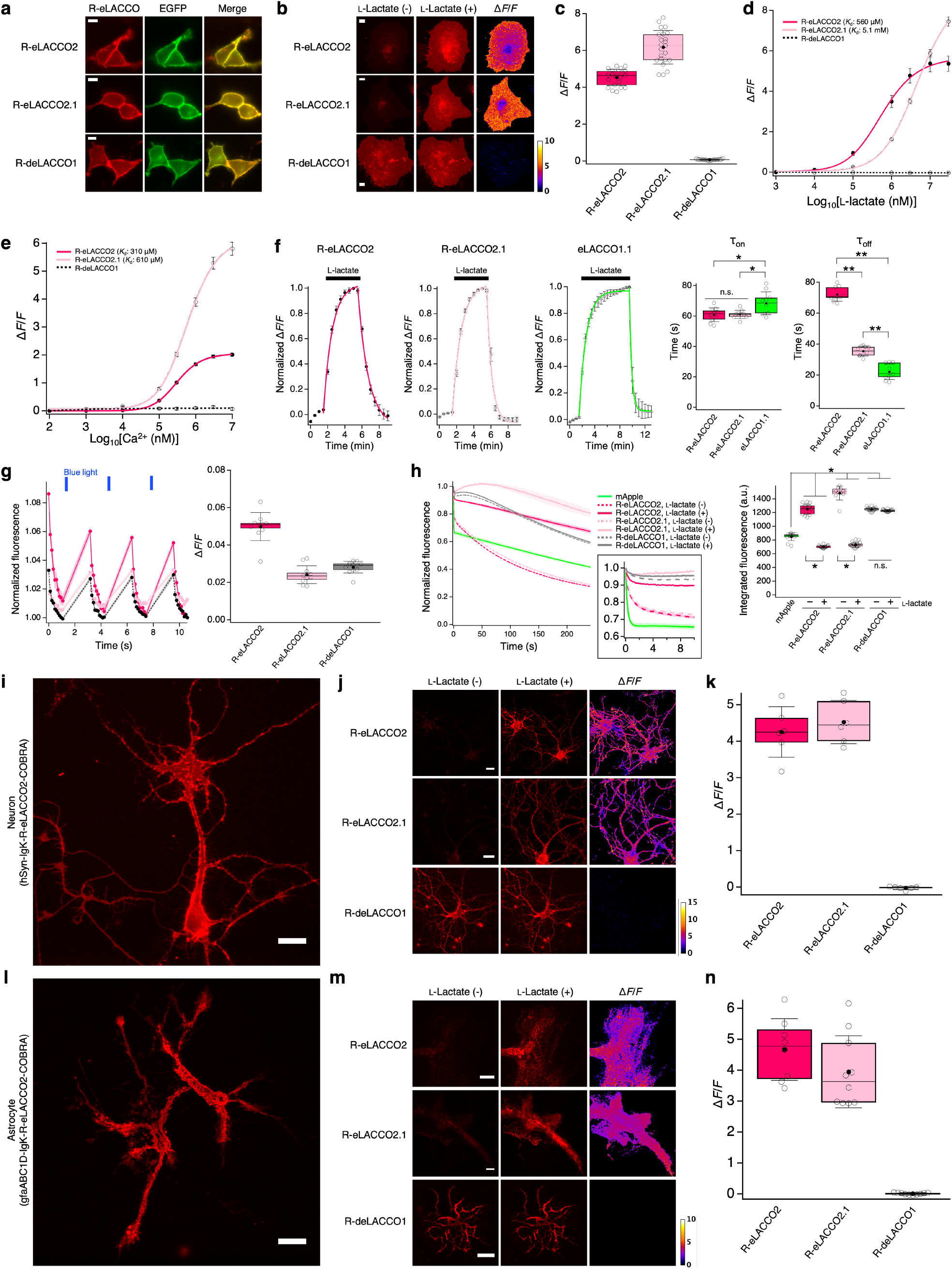
Characterization of R-eLACCO2 variants in live mammalian cells. (**a**) Localization of high-affinity R-eLACCO2, low-affinity R-eLACCO2.1, and control biosensor R-deLACCO1 with the optimized leader (Ig*κ*) and GPI anchor (COBRA) in HEK293T cells. EGFP with CD59 leader and CD59 GPI anchor was used as cell surface marker. Scale bars, 10 μm. (**b**) Fluorescence images of R-eLACCO variants expressed on cell surface of HeLa cells before and after 10 mM L-lactate treatment. Scale bars, 10 μm. (**c**) L-Lactate-dependent Δ*F*/*F* of R-eLACCO variants expressed on HeLa cells. *N* = 22, 23, and 19 cells for R-eLACCO2, R-eLACCO2.1, and R-deLACCO1, respectively. (**d**) *In situ* L-lactate titration of R-eLACCO variants expressed on HeLa cells. *n* = 11, 16, and 12 cells for R-eLACCO2, R-eLACCO2.1, and R-deLACCO1, respectively (mean ± s.e.m.). (**e**) *In situ* Ca^2+^ titration of R-eLACCO variants expressed on HeLa cells in the presence (10 mM) of L-lactate. *n* = 25, 18, and 13 cells for R-eLACCO2, R-eLACCO2.1, and R-deLACCO1, respectively (mean ± s.e.m.). (**f**) Time course (left) and time constant (right) of the fluorescence response of R-eLACCO2, R-eLACCO2.1, and eLACCO1.1 on the surface of HeLa cells. The plots represent mean ± s.d. and were fitted with a single exponential function. *n* = 10 cells for R-eLACCO2, R-eLACCO2.1, and eLACCO1.1, respectively. One-way ANOVA followed by Tukey’s multiple comparisons. \**p* < 0.05, \*\**p* < 0.0001. (**g**) Fluorescence traces and Δ*F*/*F* in response to blue light (460–480 nm) in HeLa cells expressing R-eLACCO variants. *n* = 11, 11, and 12 cells for R-eLACCO2, R-LACCO2.1, and R-deLACCO1, respectively. Fluorescence traces represent mean ± s.e.m. (**h**) Photobleaching curves (left) and integrated fluorescence (right) of R-eLACCO variants in the presence (10 mM) and absence of L-lactate and mApple expressed on the surface of HeLa cells using one-photon wide-field microscopy. Inset shows the photobleaching curves in the first 10 seconds. *n* = 12, 16, 16, 10, 18, 17, and 17 cells for mApple, R-eLACCO2 (L-lactate -/+), R-eLACCO2.1 (L-lactate -/+), and R-deLACCO1 (L-lactate -/+), respectively. In the photobleaching curves, solid lines represent mean value and shaded area represent s.e.m. One-way ANOVA followed by Tukey’s multiple comparison. \**p* < 0.0001. (**i**) Fluorescence images of primary cultured rat cortical neurons expressing R-eLACCO2. Scale bar, 20 μm. (**j**) Fluorescence images of R-eLACCO variants expressed on cell surface of rat cortical neurons before and after 10 mM L-lactate stimulation. Scale bars, 20 μm. (**k**) L-Lactate-dependent Δ*F*/*F* of R-eLACCO variants expressed on primary neurons. *n* > 10 neurons over 6 wells per construct. (**l**) Fluorescence images of primary cultured rat cortical astrocytes expressing R-eLACCO2. Scale bar, 20 μm. (**m**) Fluorescence images of R-eLACCO variants expressed on cell surface of rat cortical astrocytes before and after 10 mM L-lactate stimulation. Scale bars, 20 μm. (**n**) L-Lactate-dependent Δ*F*/*F* of R-eLACCO variants expressed on primary astrocytes. *n* = 8, 8, and 10 cells for R-eLACCO2, R-LACCO2.1, and R-deLACCO1, respectively. In the box plots of (c), (f), (g), (h), (k), and (n), the horizontal line is the median; the top and bottom horizontal lines are the 25^th^ and 75^th^ percentiles for the data; and the whiskers extend one standard deviation range from the mean represented as black filled circle.

Many cpmApple-based biosensors can undergo photoactivation when illuminated with blue light^38^, hampering their combined use with blue light-responsive biosensors and optogenetic tools such as GCaMP^24^ and channelrhodopsin (ChR)^39^. In addition, RFPs display a complexed photostability behavior compared to GFP^40^. Accordingly, we undertook a cell-based characterization of R-eLACCO variants in terms of photostability and the possibility of photoactivation. We tested the R-eLACCO variants for photoactivation upon blue-light illumination (∼4 mW cm^-2^ at 470 nm) in HeLa cells, and found that blue-light illumination elicited a small increase in fluorescence intensity of the R-eLACCO variants (**Fig. 4g**). This result indicates that the photoactivation and recovery of the R-eLACCO variants occurs with much faster kinetics and with far smaller Δ*F*/*F* (∼0.02−0.05) than L-lactate dependent fluorescence response (**Fig. 4c,d,f**). To test photostability, we continuously illuminated R-eLACCO-expressing HeLa cells using one-photon wide-field microscopy (∼4 mW cm^-2^ at 560 nm, **Fig. 4h**). R-eLACCO2 and R-eLACCO2.1 showed higher photostability (integrated fluorescence, IF, of 1251 ± 18 and 1485 ± 32, mean ± s.d., respectively) than the parent mApple RFP (IF of 848 ± 17, mean ± s.d.) in the absence of L-lactate, and lower photostability (IF of 702 ± 5 for R-eLACCO2 and 731 ± 9 for R-eLACCO2.1, mean ± s.d.) in the presence of L-lactate. R-deLACCO1 was more photostable than mApple in both the presence (IF of 1229 ± 4, mean ± s.d.) and absence (IF of 1250 ± 7, mean ± s.d.) of L-lactate.

### Characterization of R-eLACCO2 variants in neural cells

The ANLS hypothesis states that astrocytes can release L-lactate into the extracellular space and that this L-lactate is taken up by neurons to serve as an energy source^9,10^. To characterize the performance of R-eLACCO variants on the surface of neurons, we expressed them under the control of a neuron-specific promoter human synapsin (hSyn) in rat primary cortical neurons. We observed that neurons expressing R-eLACCO2 exhibited bright membrane-localized fluorescence (**Fig. 4i**). Upon bath application of 10 mM L-lactate, R-eLACCO2 and R-eLACCO2.1 had Δ*F*/*F* values of 4.3 ± 0.3 and 4.5 ± 0.2 (mean ± s.e.m.), respectively, whereas R-deLACCO1 had no response (**Fig. 4j,k**). To determine whether R-eLACCO variants could sense the extracellular L-lactate on the surface of astrocytes, we expressed them under the control of an astrocyte-specific promoter gfaABC1D in rat primary cortical astrocytes. The fluorescence imaging revealed that the biosensors were expressed (**Fig. 4l**) and R-eLACCO2 and R-eLACCO2.1 displayed Δ*F*/*F* values of 4.7 ± 0.4 and 3.9 ± 0.4 (mean ± s.e.m., **Fig. 4m,n**), respectively, upon bath application of 10 mM L-lactate. R-deLACCO1 showed no response to L-lactate, similar to our results obtained with HeLa cells and neurons. Taken together, these results indicated that R-eLACCO variants with the optimized leader and anchor can be used for imaging of extracellular L-lactate concentration dynamics on the surfaces of neurons and astrocytes.

## Discussion

This study describes the development of a red fluorescent genetically encoded extracellular L-lactate biosensor, designated R-eLACCO2. Swapping the GFP-derived domain of eLACCO1.1 for cpmApple led to a biosensor prototype in which the L-lactate-dependent conformational change of TTHA0766 allosterically modulates the red fluorescence of cpmApple. Extensive directed evolution and site-directed mutagenesis of the prototype led to the high-performance L-lactate biosensor, R-eLACCO2. Structure-based rational mutagenesis provided the R-eLACCO2.1 variant with optimal L-lactate affinity for the physiological concentration range of extracellular L-lactate. An intensive effort to optimize the combination of leader and anchor domain for targeting the affinity-tuned variant to the extracellular environment identified the Ig*κ*-derived N-terminal leader sequence and the COBRA-based C-terminal anchor, as the optimal combination for imaging of extracellular L-lactate.

To the best of our knowledge, R-eLACCO2 has the highest fluorescence response of any genetically encoded red fluorescent biosensor as measured with purified protein and in the extracellular environment of mammalian cells (**Table S1**)^22^. However, there are several important considerations that users should pay close attention to. As with eLACCO1, the TTHA0766-based R-eLACCO2 only functions as an L-lactate biosensor in the presence of Ca^2+^. For extracellular applications this should not be a major concern because the apparent *K*_d_s of R-eLACCO2 and R-eLACCO2.1 for Ca^2+^ are substantially lower than the physiological and pathological Ca^2+^ concentration range (∼1.5−1.7 mM) in the extracellular space of brain tissue (**Fig. 4e**)^37^. However, given that the Ca^2+^ concentration in the extracellular environment can transiently decrease to ∼1 mM during neural activity artificially evoked by extreme stimulations such as long trains of action potentials^41^, careful consideration should be taken when interpreting the fluorescence signal of R-eLACCO2 variants in brain tissue. A change in extracellular Ca^2+^ concentration from 1.7 mM to 1.0 mM could cause up to a -8.6% or -15% fluorescence intensity change for R-eLACCO2 and R-eLACCO2.1 respectively (**Fig. 4e**). Another important consideration for the application of R-eLACCO2 is the kinetics of fluorescence response. R-eLACCO2 variants, similar to eLACCO1.1, show relatively slow kinetics of fluorescence response compared to GCaMP^24^ and R-GECO1 (ref. 31) (**Fig. 4f**). Previous work using enzyme-mediated electrodes has revealed that the concentration of L-lactate in the extracellular space typically changes on the time scale of minutes in brain tissue^16^. As lactate dynamics are much slower than neural activity-dependent intracellular Ca^2+^ dynamics (∼ ms to s), R-eLACCO2 variants are likely to be well-suited for investigating physiologically relevant changes in the extracellular L-lactate concentration in brain tissue.

The emerging roles of intercellular shuttling of L-lactate have highlighted the need for the development of genetically encoded biosensors for extracellular L-lactate. This need previously motivated us to engineer a green fluorescent extracellular L-lactate biosensor eLACCO1.1 (ref. 29). However, studies of metabolism will likely require simultaneous visualization of multiple other metabolites such as glucose and ATP and/or visualization of L-lactate in different cell types or different regions of single cells. To date, the most effective and widely used biosensors and optogenetic actuators require a blue-light excitation^22,30^, hindering multiplexed imaging of extracellular L-lactate with other targets. To help to surmount this obstacle, we have now expanded the color spectrum of eLACCO1.1 by engineering a red fluorescent biosensor, R-eLACCO2. To the best of our knowledge, no red fluorescent biosensors for extracellular or intracellular^25−28^ L-lactate have been reported. R-eLACCO2 is a first-in-class red fluorescent biosensor for minimally-invasive imaging of the dynamics of extracellular L-lactate, potentially enabling an orthogonal readout of distinct and multiple metabolic events.

## Methods

### General methods and materials

The gene encoding the lactate binding bacterial periplasmic protein TTHA0766 was derived from the gene encoding the previously reported L-lactate biosensor eLACCO1 (ref. 29). The gene encoding cpmApple was derived from the gene encoding the previously reported Ca^2+^ biosensor R-GECO1 (ref. 31). Phusion high-fidelity DNA polymerase (Thermo Fisher Scientific) was used for routine polymerase chain reaction (PCR) amplifications, and Taq DNA polymerase (New England Biolabs) was used for error-prone PCR. Q5 high-fidelity DNA polymerase (New England Biolabs) was used for site-directed mutagenesis^42^. Restriction endonucleases, rapid DNA ligation kits, and GeneJET miniprep kits were purchased from Thermo Fisher Scientific. PCR products and products of restriction digests were purified using agarose gel electrophoresis and the GeneJET gel extraction kit (Thermo Fisher Scientific). DNA sequences were analyzed by DNA sequence service of Fasmac Co., Ltd. Fluorescence excitation and emission spectra were recorded on a Spark plate reader (Tecan).

### Engineering of R-eLACCO2

The gene encoding cpmApple with N- and C-terminal linkers (DW and EADG, respectively) was amplified using the R-GECO1 gene as template, followed by insertion between the codons encoding Ile191 and Gly192 of TTHA0766 in pBAD-eLACCO1 (ref. 29) by Gibson assembly (New England Biolabs). The resulting variant, designated R-eLACCO0.1, was subjected to an iterative process of library generation and screening in *E. coli* strain DH10B (Thermo Fisher Scientific) in LB media supplemented with 100 μg mL^-1^ ampicillin and 0.02% L-arabinose. Libraries were generated by site-directed mutagenesis or error-prone PCR of the whole gene. In the screening, proteins were extracted using B-PER bacterial protein extraction reagent (Thermo Fisher Scientific) and tested for fluorescence brightness and lactate-dependent response. For each round, approximately 100−400 fluorescent colonies were picked, cultured and tested on 96-well plates under a plate reader. There were 22 rounds of screening and one site-directed mutagenesis (Ile191Val) before R-eLACCO2 was identified. Leu79Ile mutation was added to R-eLACCO2 to tune the lactate affinity by extending the R-eLACCO2 template using Q5 high-fidelity DNA polymerase and oligonucleotides containing Leu79Ile mutation (CTG to ATC) followed by digestion with DpnI (Thermo Fisher Scientific). The resulting low-affinity mutant was designated as R-eLACCO2.1. Asp441Asn mutation was added to R-eLACCO2 to abrogate the lactate affinity. Six rounds of directed evolution were performed to improve the brightness of R-eLACCO2 Asp441Asn. The resulting mutant was designated as R-deLACCO1.

### Protein purification and *in vitro* characterization

The gene encoding R-eLACCO variants, with a poly-histidine tag on the N-terminus, was expressed from the pBAD vector. Bacteria were lysed with a sonicator (Branson) and then centrifuged at 15,000*g* for 20 min, and proteins were purified by Ni-NTA affinity chromatography (Agarose Bead Technologies). Absorption spectra of the samples were collected with a Lambda950 Spectrophotometer (PerkinElmer) and UV-1800 Spectrophotometer (Shimadzu). To perform pH titrations, protein solutions were diluted into buffers (pH from 4 to 11) containing 30 mM trisodium citrate, 30 mM sodium borate, 30 mM MOPS, 100 mM KCl, 1 mM CaCl_2_, and either no L-lactate or 100 mM L-lactate. Fluorescence intensities as a function of pH were then fitted by a sigmoidal binding function to determine the p*K*_a_. For lactate titration, buffers were prepared by mixing an L-lactate (-) buffer (30 mM MOPS, 100 mM KCl, 1 mM CaCl_2_, pH 7.2) and an L-lactate (+) buffer (30 mM MOPS, 100 mM KCl, 1 mM CaCl_2_, 100 mM L-lactate, pH 7.2) to provide L-lactate concentrations ranging from 0 mM to 100 mM at 25 °C. Fluorescence intensities were plotted against L-lactate concentrations and fitted by a sigmoidal binding function to determine the Hill coefficient and apparent *K*_d_. For Ca^2+^ titration, buffers were prepared by mixing a Ca^2+^ (-) buffer (30 mM MOPS, 100 mM KCl, 10 mM EGTA, 100 mM L-lactate, pH 7.2) and a Ca^2+^ (+) buffer (30 mM MOPS, 100 mM KCl, 10 mM CaEGTA, 100 mM L-lactate, pH 7.2) to provide Ca^2+^ concentrations ranging from 0 μM to 39 μM at 25 °C. Buffers with Ca^2+^ concentrations more than 39 μM was prepared by mixing a Ca^2+^ (-, w/o EGTA) buffer (30 mM MOPS, 100 mM KCl, 100 mM L-lactate, pH 7.2) and a Ca^2+^ (+, w/o EGTA) buffer (30 mM MOPS, 100 mM KCl, 100 mM CaCl_2_, 100 mM L-lactate, pH 7.2).

To collect the two-photon absorption spectra, the tunable femtosecond laser InSight DeepSee (Spectra-Physics, Santa Clara, CA) was used to excite the fluorescence of the sample contained within a PC1 Spectrofluorometer (ISS, Champaign, IL). The details of the method and protocols are described in ref. 43 To measure the two-photon excitation spectral shapes, we used short-pass filters 633SP and 770SP in the emission channel. LDS 798 and Coumarin 540A were used as spectral standards. Quadratic power dependence of fluorescence intensity in the proteins and standards was checked at several wavelengths across the spectrum.

The two-photon cross section (σ_2_) of the anionic form of the chromophore was measured as described previously^44^. Rhodamine 6G in MeOH was used as a reference standard with excitation at 1060 nm (ref. 43). For the L-lactate free state, the σ_2_ of the neutral form of the chromophore was also measured versus fluorescein (Millipore Sigma, Darmstadt, Germany) in 10 mM NaOH (pH 12) at 820 and 840 nm (ref. 43). Extinction coefficients were determined by alkaline denaturation as detailed in ref. 35. The two-photon absorption spectra were normalized to the measured σ_2_ values. To normalize to the total two-photon brightness (*F*_2_), the spectra were then multiplied by the quantum yield and the relative fraction of the respective form of the chromophore for which the σ_2_ was measured. The data is presented this way because R-eLACCO1.1 and R-eLACCO2 contain a mixture of the neutral and anionic forms of the cpmApple chromophore. This is described in further detail in refs. 35 and 45.

### Construction of mammalian expression vectors

For cell surface expression, the genes encoding R-eLACCO2, R-eLACCO2.1, and R-deLACCO1 were amplified by PCR followed by digestion with BglII and EcoRI, and then ligated into a pcDNA3.1 vector (Thermo Fisher Scientific) that contains N-terminal leader sequence and C-terminal anchor. To construct CD59-anchored R-eLACCO2.1 with various leader sequences, complementary oligonucleotides (Thermo Fisher Scientific) encoding each leader sequence were digested by XhoI and BglII, and then ligated into a similarly-digested pcDNA3.1 including CD59-anchored R-eLACCO2.1. The gene encoding PDGFR transmembrane domain was amplified by PCR using pDisplay vector (Thermo Fisher Scientific) as a template, and then substituted with CD59 GPI domain of the pcDNA3.1 product above by using EcoRI and HindIII. To construct R-eLACCO plasmids for expression in neurons and astrocytes, the gene encoding R-eLACCO variants including the Ig*κ* leader and COBRA GPI anchor sequence in the pcDNA vector was first amplified by PCR followed by digestion with NheI and XhoI, and then ligated into pAAV plasmid containing the hSyn and gfaABC1D promoter.

### Imaging of R-eLACCO variants in HEK293T and HeLa cell lines

HEK293T and HeLa cells were maintained in Dulbecco’s modified Eagle medium (DMEM; Nakalai Tesque) supplemented with 10% fetal bovine serum (FBS; Sigma-Aldrich) and 1% penicillin-streptomycin (Nakalai Tesque) at 37 °C and 5% CO_2_. Cells were transiently transfected with the constructed plasmid using polyethyleneimine (Polysciences). Transfected cells were imaged using a IX83 wide-field fluorescence microscopy (Olympus) equipped with a pE-300 LED light source (CoolLED), a 40× objective lens (numerical aperture (NA) = 1.3; oil), an ImagEM X2 EM-CCD camera (Hamamatsu), Cellsens software (Olympus), and a STR stage incubator (Tokai Hit). The filter sets used in live cell imaging had the following specification. eLACCO1.1 and EGFP: excitation 470/20 nm, dichroic mirror 490-nm dclp, and emission 518/45 nm; R-eLACCO variants: excitation 545/20 nm, dichroic mirror 565-nm dclp, and emission 598/55 nm. Fluorescence images were analyzed with ImageJ software (National Institutes of Health).

For the optimization of leader and anchor, HEK293T cells were co-transfected with respective R-eLACCO2.1 genes and pAMEXT-EGFP. Measurement of individual fluorescence distribution profiles and calculation of Pearson correlation coefficient were carried out as follows: linear region of interest (ROI) was generated on the cell of interest to cross the membrane manually. The fluorescence distribution of ROI was measured from both R-eLACCO stack and GFP marker stack using the Plot Profile function (ImageJ2 V2.3.0/1.53q). After measuring the raw values, the fluorescence profile data from both stacks were combined by means of a home-made python script. Pandas (version 0.25.1) python package was used for handling data as well as computing descriptive statistics. The correlation function used in this analysis was the Pearson Correlation score expressed by the *corr* function in Pandas.

For L-lactate bath application to measure *in situ K*_d_ and response kinetics, HeLa cells seeded onto coverslips were transfected with R-eLACCO variants or eLACCO1.1 (ref. 29). Forty-eight hours after transfection, the transfected HeLa cells were washed twice with Hank’s balanced salt solution (HBSS(+); Nakalai Tesque), and then the coverslips were transferred into Attofluor™ Cell Chamber (Thermo Fisher Scientific, Cat. #A7816) with HBSS(+) supplemented with 10 mM HEPES (Nakalai Tesque) and 1 μM AR-C155858 (Wako). Rapid change of bath solutions during the image was performed in a remove-and-add manner using a homemade solution remover^46^. For photostability test, R-eLACCO variants or mApple was expressed on the surface of HeLa cells and illuminated by excitation light at 100% intensity of LED (∼4 mW cm^-2^ on the objective lens), and then their fluorescence images were recorded at 37 °C for 4 min with the exposure time of 50 ms and no interval time. For blue-light-mediated photoactivation, HeLa cells transfected with R-eLACCO variants were imaged under intermittent illumination (exposure time for 200 ms per illumination) of LED light filtered with 470/20 bandpath (∼4 mW cm^-2^ on the objective lens).

For imaging of Ca^2+^-dependent fluorescence, HeLa cells seeded onto coverslips were transfected with R-eLACCO variants. Forty-eight hours after transfection, the coverslips were transferred into Attofluor™ Cell Chamber with HBSS(-) buffer (Nakalai Tesque) supplemented with 10 mM HEPES and 10 mM L-lactate. Other bath solutions were supplemented with Ca^2+^ of 100 nM, 1 μM, 10 μM, 100 μM, 300 μM, 1 mM, 3 mM, and 10 mM. Rapid change of bath solutions during the image was performed in a remove-and-add manner using a homemade solution remover^46^.

### Imaging of R-eLACCO variants in primary neurons and astrocytes

Male and female pups were obtained from a single timed-pregnant Sprague Dawley rat (Charles River Laboratories, purchased from Japan SLC, Inc.). Experiments were performed with cortical/hippocampal primary cultures from the E21 (after C-section of the pregnant rat) plated in glass-bottom 24-well plates (Cellvis) where 0.5 × 10^6^ cells were used for three wells. Cultures were nucleofected at time of plating with Nucleofector 4D (Lonza), and imaged 14 days later. Three wells were plated and imaged per nucleofected construct. Neurons were cultured in NbActive4 medium (BrainBits) supplemented with 1% penicillin-streptomycin at 37 °C and 5% CO_2_. Astrocytes were cultured in DMEM supplemented with 10% FBS and 1% penicillin-streptomycin at 37 °C and 5% CO_2_. Culture media were replaced with 1 mL of imaging buffer (145 mM NaCl, 2.5 mM KCl, 10 mM glucose, 10 mM HEPES, 2 mM CaCl_2_, 1 mM MgCl_2_, pH 7.4) for imaging^24^.

### Animal care

For experiments performed at The University of Tokyo, all methods for animal care and use were approved by the institutional review committees of School of Science, The University of Tokyo.

### Statistics and reproducibility

All data are expressed as mean ± s.d. or mean ± s.e.m., as specified in figure legends. Box plots are used for Figs. 3b,d and 4c,f,g,h,k,n. In these box plots, the horizontal line is the median; the top and bottom horizontal lines are the 25^th^ and 75^th^ percentiles for the data; and the whiskers extend one standard deviation range from the mean represented as black filled circle. Sample sizes (*n*) are listed with each experiment. No samples were excluded from analysis and all experiments were reproducible. For pharmacological specificity, statistical analysis was performed using one-way analysis of variance (ANOVA) with the Dunnett’s post hoc tests (Igor Pro 8). In *in situ* kinetics analysis and photobleaching experiments, group differences were analyzed using one-way ANOVA followed by Tukey’s multiple comparison (GraphPad Prism 9). Microsoft Excel software was used to plot for Figs. 1c,d and 2b,d,g.

## Data availability

The data and plasmids encoding R-eLACCO variants that support the findings of this study are available from the corresponding authors on reasonable request.

## Acknowledgments

The authors thank M. Yamane, Z. Wenchao, A. Aggarwal, and T. Terai for technical support. Work at the University of Tokyo was supported by the Japan Society for the Promotion of Science (Grants-in-Aid for Early-Career Scientists 19K15691 and 21K14738, Grant-in-Aid for Scientific Research A 19H01015, and Grants-in-Aid for Scientific Research S 19H05633), Toyota Physical and Chemical Research Institute, The Precise Measurement Technology Promotion Foundation, and Strategic Research Program for Brain Sciences from the Japan Agency for Medical Research and Development (JP21dm0207073). Work at Montana State University was supported by National Institutes of Health (NIH) grants U01 NS094246, U24 NS109107, and F31 NS108593.

## Author Contributions

Y.N. developed all R-eLACCO variants and performed characterization *in vitro* and in mammalian cells. Y.N. and Y.K. performed screening of leader and anchor sequences, and cell imaging. R.H. purified R-eLACCO1.1 protein for two-photon characterization. M.D. measured two-photon excitation spectra, extinction coefficient, and quantum yield. H.S., Y.H., and T.T. supervised imaging of rat primary cells. Y.N. and R.E.C. supervised research. Y.N. and R.E.C. wrote the manuscript.

## Supplementary Figures

**Supplementary Figure 1.**
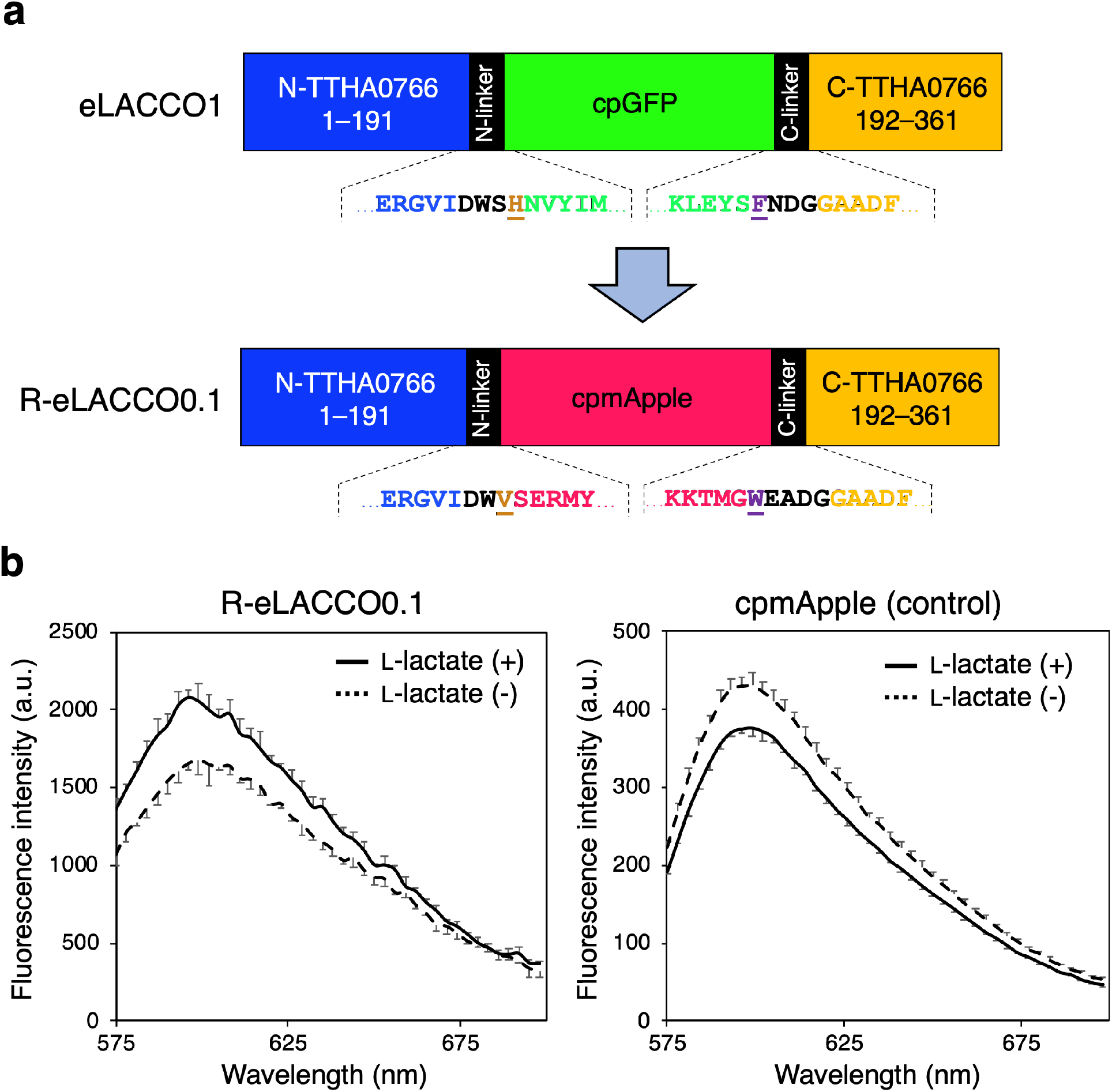
Construction of the biosensor prototype. (**a**) Schematic illustration of the biosensor prototype construction. Linker regions are shown in black and the two “gate post” residues^1^ in cpGFP or cpmApple are highlighted in dark orange and purple. (**b**) Emission spectra of R-eLACCO0.1 and cpmApple in the presence (10 mM) and absence of L-lactate. Error bars represent standard deviation of triplicates.

**Supplementary Figure 2.**
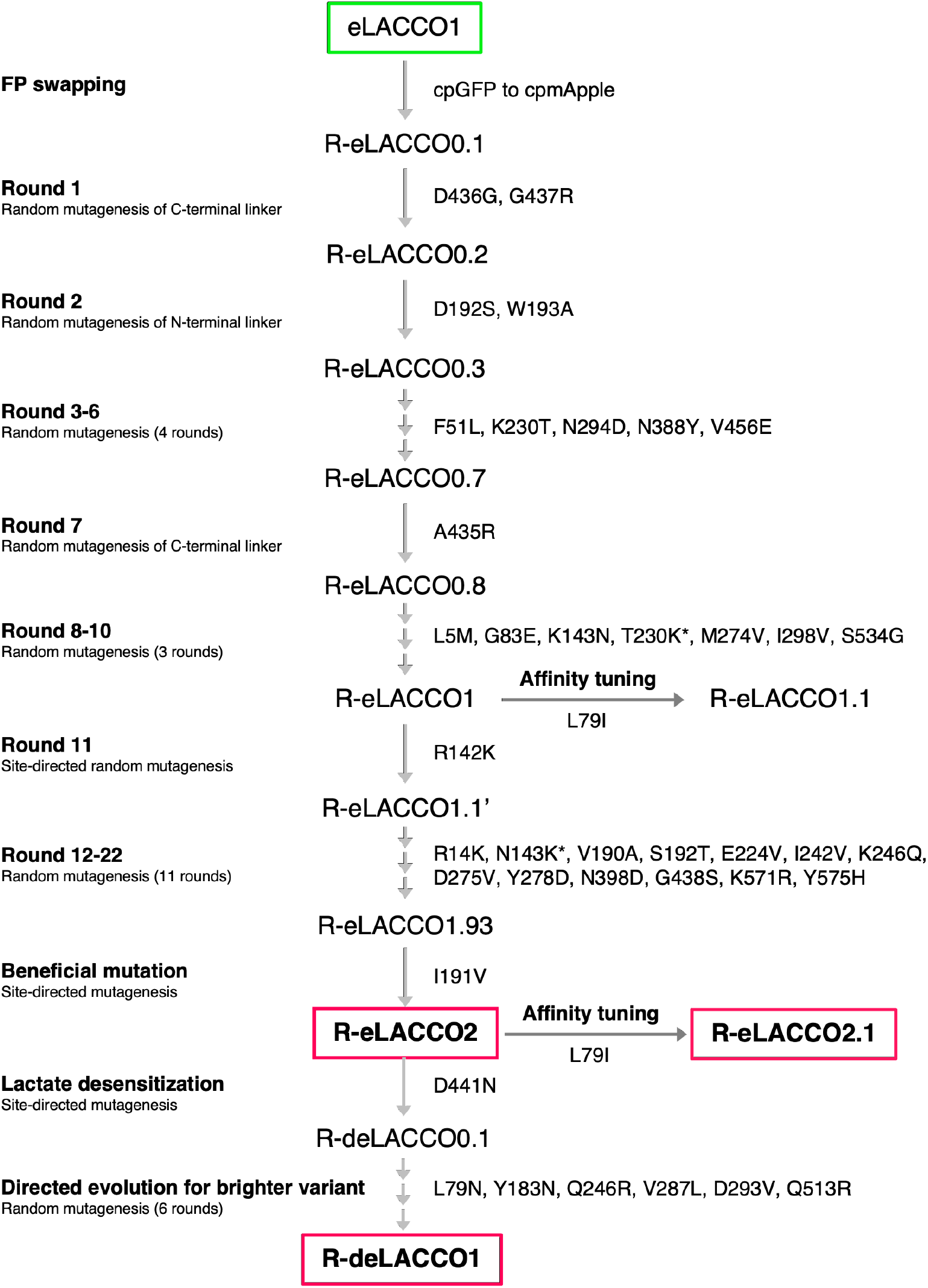
Lineage of R-eLACCO variants from eLACCO1 (ref. 2). Asterisks represent reversions.

**Supplementary Figure 3.**
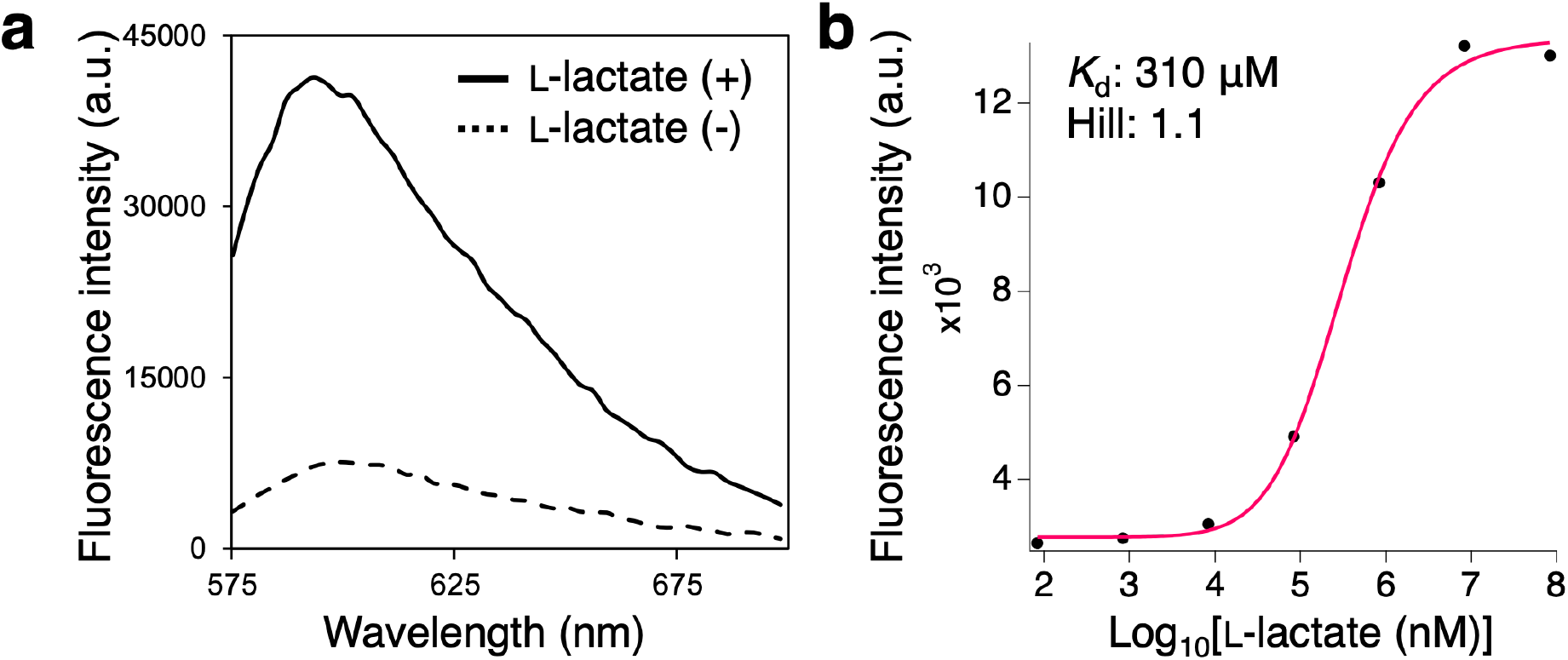
*In vitro* characterization of R-eLACCO1. (**a**) Emission spectra of R-eLACCO1 in the presence (10 mM) and absence of L-lactate. (**b**) Dose-response curve of R-eLACCO1 for L-lactate.

**Supplementary Figure 4.**
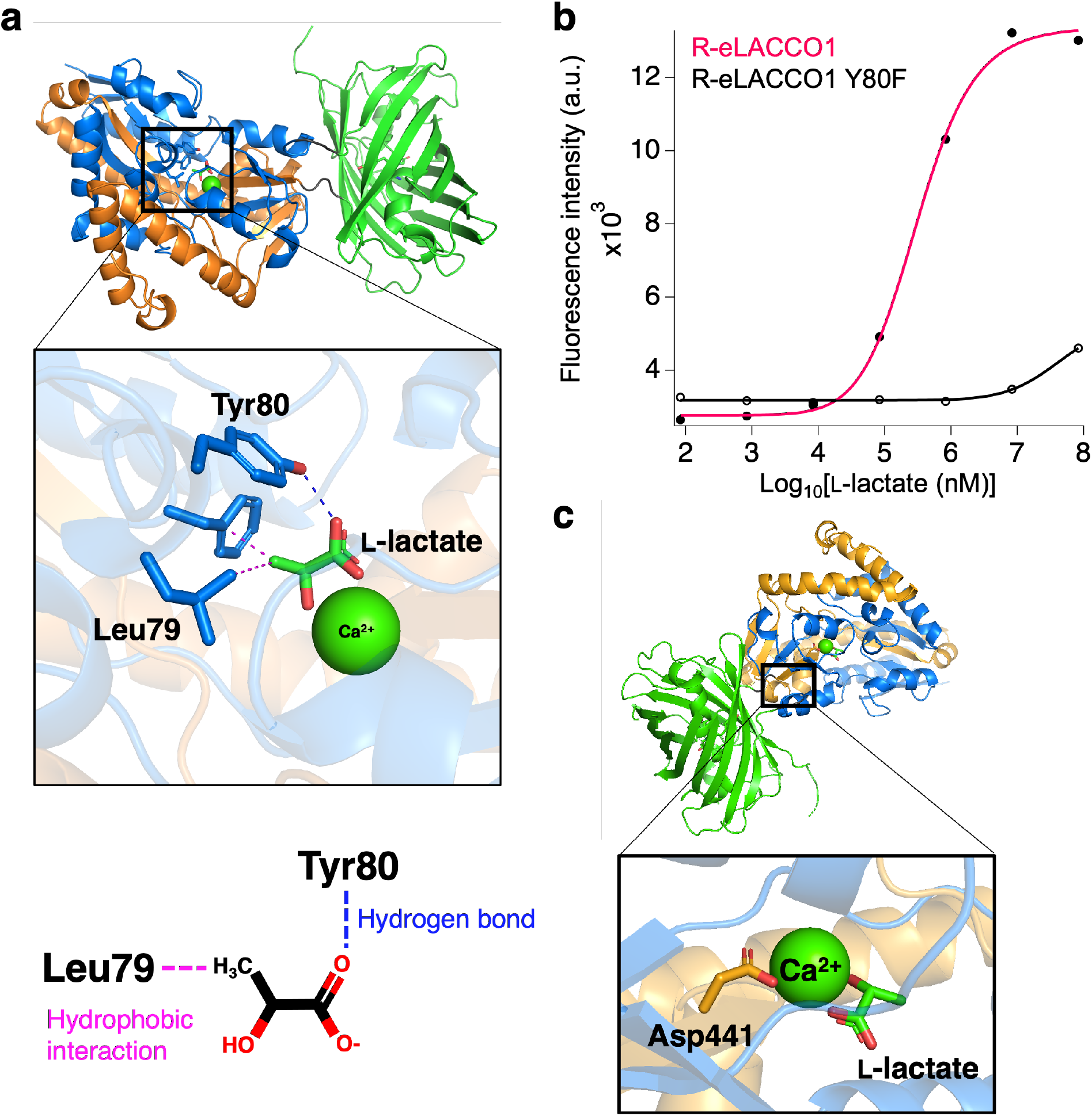
Affinity tuning of R-eLACCO1. (**a**) Crystal structure of eLACCO1 (PDB 7E9Y) and zoom-in view of the L-lactate binding pocket. The phenol group of the Tyr80 side chain forms a hydrogen bond to the carboxylate group of L-lactate. The hydrophobic side chain of Leu79 interacts with the methyl group of L-lactate. (**b**) Fluorescence of R-eLACCO1 and its Tyr80Phe variant as a function of L-lactate. (**c**) Crystal structure of eLACCO1 and zoom-in view of the Ca^2+^ binding pocket. Carboxyl group of side chain of Asp441 (R-eLACCO numbering) coordinates to Ca^2+^.

**Supplementary Figure 5.**
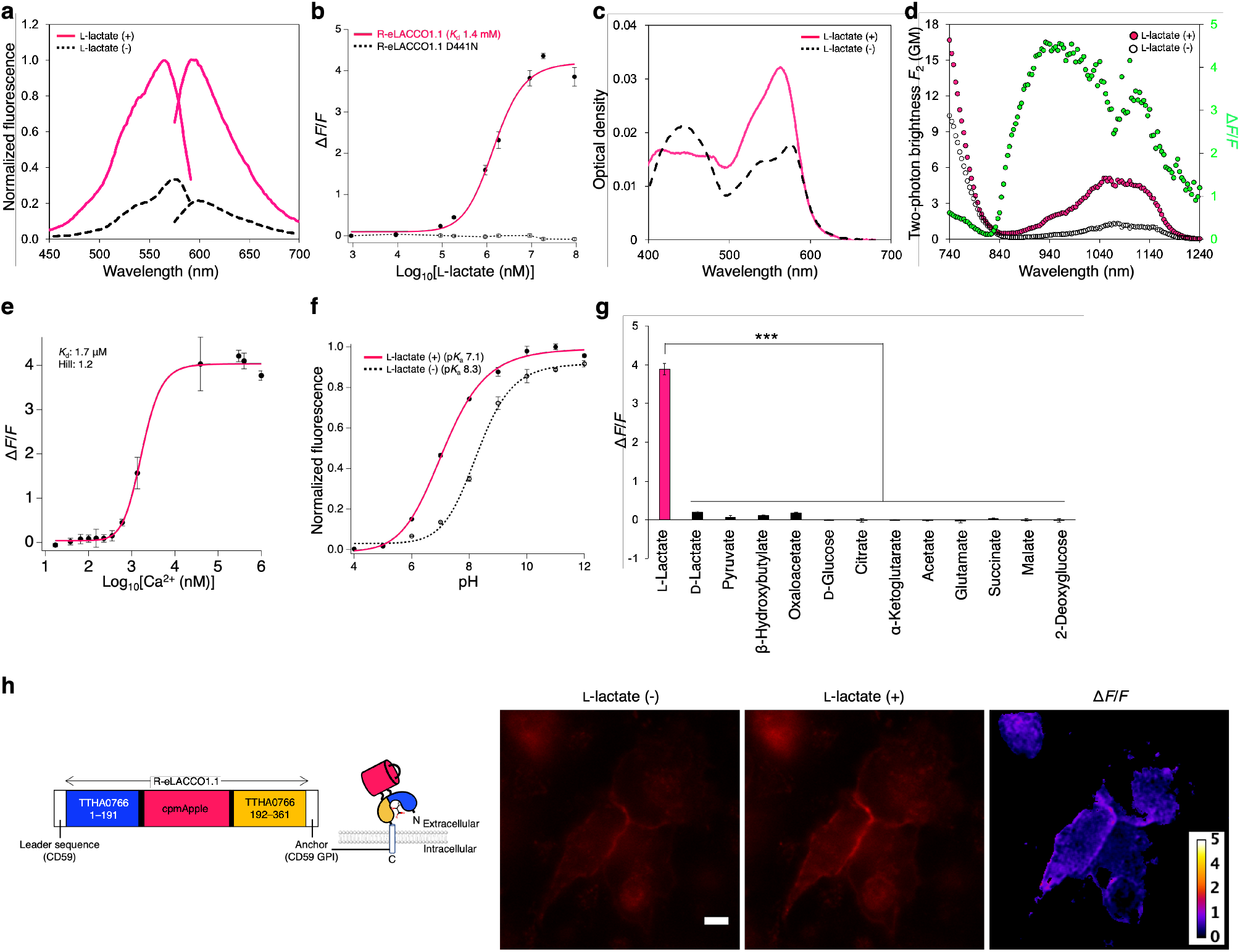
Characterization of R-eLACCO1.1. (**a**) Excitation and emission spectra of R-eLACCO1.1 in the presence (10 mM) and absence of L-lactate. (**b**) Dose-response curves of R-eLACCO1.1 and its D441N variant for L-lactate. *n* = 3 experimental triplicates (mean ± s.d.). (**c**) Absorbance spectra of R-eLACCO1.1 in the presence (10 mM) and absence of L-lactate. (**d**) Two-photon excitation spectra of R-eLACCO1.1 in the presence (10 mM) and absence of L-lactate. Δ*F*/*F* is represented in the green plots. GM, Goeppert-Mayer units. (**e**) Dose-response curve of R-eLACCO1.1 as a function of Ca^2+^ in the presence (10 mM) of L-lactate. *n* = 3 experimental triplicates (mean ± s.d.). (**f**) pH titration curves of R-eLACCO1.1 in the presence (10 mM) and absence of L-lactate. *n* = 3 experimental triplicates (mean ± s.d.). (**g**) Pharmacological specificity of R-eLACCO1.1. Concentration of each metabolite is 10 mM. *n* = 3 experimental triplicates (mean ± s.d.). Statistical analysis was performed using one-way analysis of variance (ANOVA) with the Dunnett’s post hoc tests. \*\*\**p* < 0.000001. (**h**) Fluorescence images of HeLa cells expressing R-eLACCO1.1 on cell surface before and after 10 mM L-lactate stimulation. Scale bar, 10 μm. **M F S**

**Supplementary Figure 6.**
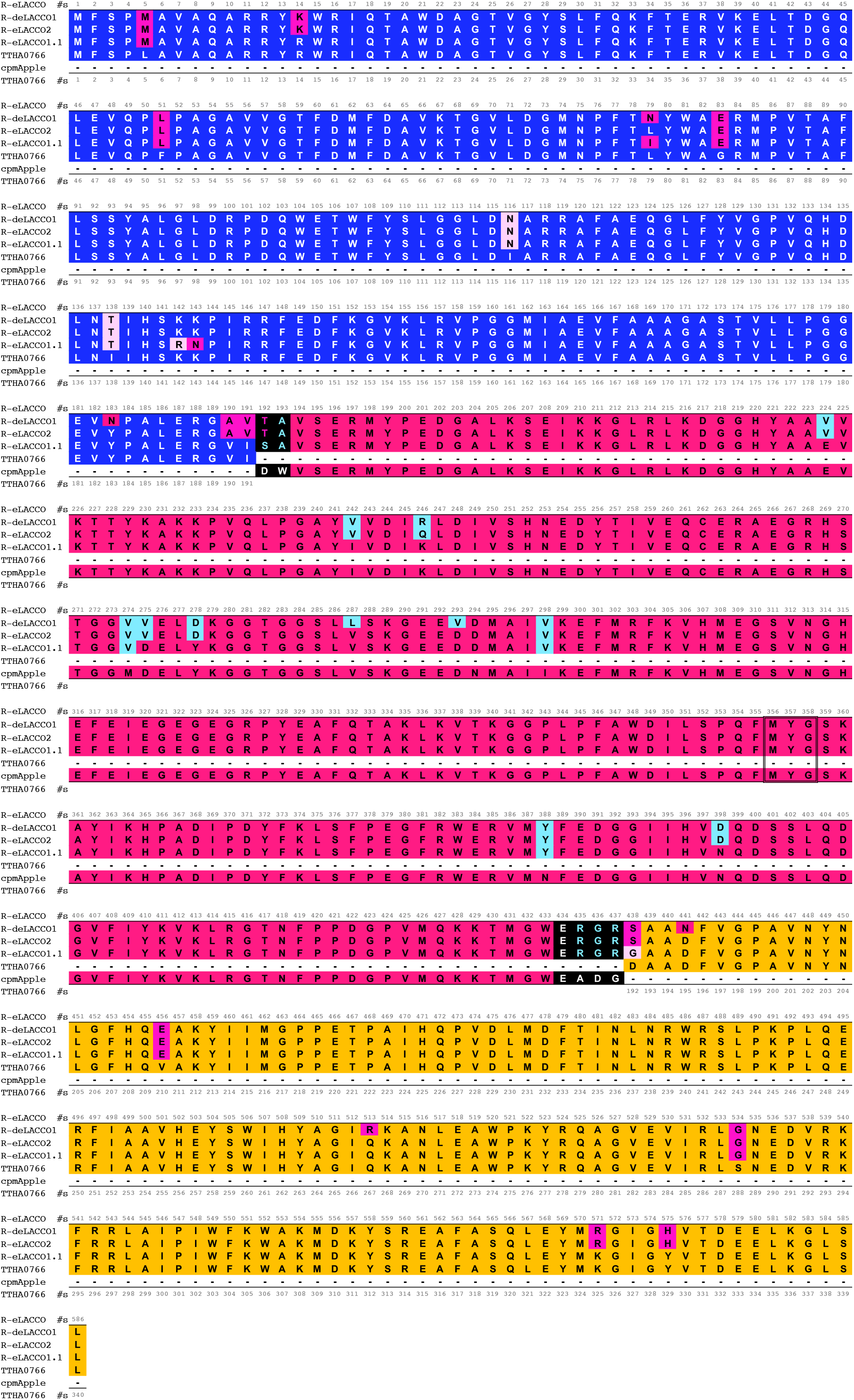
Sequence alignment of TTHA0766, cpmApple, R-eLACCO1.1, R-eLACCO2, and R-deLACCO1. Mutations in R-eLACCO1.1 and R-eLACCO2, relative to TTHA0766 and cpmApple, are highlighted in magenta and blue, respectively. Light magenta represent mutations derived from eLACCO1 (ref. 2). The chromophore-forming residues are surrounded by double line.

**Supplementary Figure 7.**
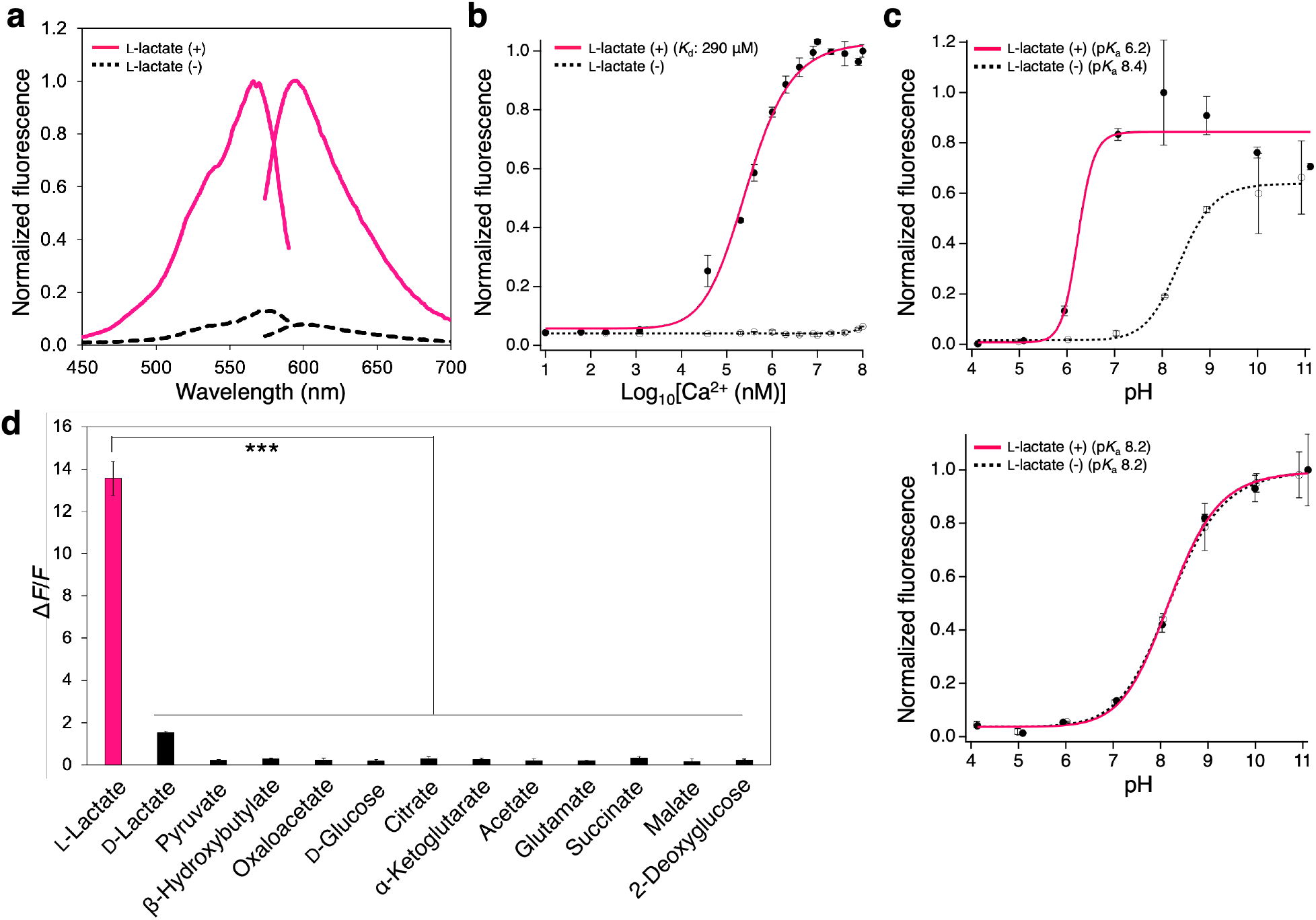
*In vitro* characterization of R-eLACCO2.1 and R-deLACCO1. (**a**) Excitation and emission spectra of R-eLACCO2.1 in the presence (10 mM) and absence of L-lactate. (**b**) Dose-response curve of R-eLACCO2.1 as a function of Ca^2+^ in the presence (100 mM) and absence of L-lactate. *n* = 3 experimental triplicates (mean ± s.d.). (**c**) pH titration curves of R-eLACCO2.1 (upper) and R-deLACCO1 (bottom) in the presence (100 mM) and absence of L-lactate. *n* = 3 experimental triplicates (mean ± s.d.). (**d**) Pharmacological specificity of R-eLACCO2.1. Concentration of each metabolite is 10 mM. *n* = 3 experimental triplicates (mean ± s.d.). Statistical analysis was performed using one-way ANOVA with the Dunnett’s post hoc tests. \*\*\**p* < 0.000001.

**Supplementary Table 1.** Biochemical parameters of R-eLACCO variants.

|  | R-eLACCO1.1 |  | R-eLACCO2 |  | R-eLACCO2.1 |  | R-deLACCO1 |  | R-GECO1<sup>a</sup> |  | eLACCO1.1<sup>b</sup> |  |
| --- | --- | --- | --- | --- | --- | --- | --- | --- | --- | --- | --- | --- |
| L-Lactate | - | + | - | + | - | + | - | + | -<sup>c</sup> | +<sup>c</sup> | - | + |
| *In vitro* (purified protein) |  |  |  |  |  |  |  |  |  |  |  |  |
| Fluorescence excitation maximum (nm) | 577 | 565 | 578 | 566 | 578 | 566 | 578 | 574 | 577 | 561 | 493 | 493 |
| Fluorescence emission maximum (nm) | 599 | 593 | 602 | 594 | 600 | 594 | 602 | 598 | 600 | 589 | 510 | 510 |
| pK<sub>a</sub> | 8.3 | 7.1 | 8.8 | 6.1 | 8.4 | 6.2 | 8.2 | 8.2 | 8.9 | 6.6 | 9.4 | 7.1 |
| $\Delta F/F^d$ | 3.9 | | 20 | | 12 | | 0.2 | | 15 | | 3.2 | |
| K<sub>d</sub> (L-lactate) (mM) | 1.4 |  | 0.46 |  | 5.7 |  | N/A |  | N/A |  | 3.9 |  |
| Hill coefficient (L-lactate) | 1.3 |  | 1.1 |  | 1.2 |  | N/A |  | N/A |  | 0.94 |  |
| K<sub>d</sub> (Ca<sup>2+</sup>) (μM) | 1.7 |  | 7.6 |  | 290 |  | ND |  | 0.48 |  | 2.0 |  |
| Hill coefficient (Ca<sup>2+</sup>) | 1.2 |  | 1.2 |  | 1.1 |  | ND |  | 2.1 |  | 1.4 |  |
| *In situ* (in live mammalian cells) |  |  |  |  |  |  |  |  |  |  |  |  |
| $\Delta F/F$ (HeLa)<sup>d</sup> | 1 | | 4.5 | | 6.5 | | 0.0 | | 3.9 | | 3.3 | |
| $\Delta F/F$ (neuron)<sup>d</sup> | ND | | 4.3 | | 4.5 | | 0.0 | | N/A | | 2.0 | |
| $\Delta F/F$ (astrocyte)<sup>d</sup> | ND | | 4.7 | | 3.9 | | 0.0 | | N/A | | N/A | |
| K<sub>d</sub> (L-lactate) (mM) | ND |  | 0.56 |  | 5.1 |  | N/A |  | N/A |  | 1.6 |  |
| Hill coefficient (L-lactate) | ND |  | 1.1 |  | 1.2 |  | N/A |  | N/A |  | N/A |  |
| K<sub>d</sub> (Ca<sup>2+</sup>) (μM) | ND |  | ND | 310 | ND | 610 | ND |  | N/A |  | N/A |  |
| Hill coefficient (Ca<sup>2+</sup>) | ND |  | ND | 1.2 | ND | 1.2 | ND |  | N/A |  | N/A |  |

N/A, not applicable. ND, not determined.
- a. Ref. 3. - b. Ref. 2. - c. $\text{Ca}^{2+}$ . - d. 10 mM L-lactate.

